# Physiological and Genetic Drivers Underpinning Canopy Development are Associated with Durum Wheat Yield in Rainfed Environments

**DOI:** 10.1101/2021.08.21.457180

**Authors:** Yichen Kang, Shanice V. Haeften, Daniela Bustos-Korts, Stjepan Vukasovic, Sana Ullah Khan, Eric Dinglasan, Jack Christopher, Karine Chenu, Jason A. Able, Millicent R. Smith, Kai P. Voss-Fels, Andries B. Potgieter, David R. Jordan, Andrew K. Borrell, Samir Alahmad, Lee T. Hickey

## Abstract

New durum wheat (*Triticum turgidum* L. ssp. *Durum*) cultivars with improved adaptation to variable rainfall environments are required to sustain productivity in the face of climate change. Physiological traits related to canopy development underpin the production of biomass and yield, as they interact with solar radiation and affect the timing of water use throughout the growing season. This study explored the temporal canopy dynamics of durum wheat using a nested-association mapping population evaluated for longitudinal normalized difference vegetation index (NDVI) measurements. Association mapping was performed to identify quantitative trait loci (QTL) for time-point NDVI and spline-smoothed NDVI trajectory traits. Yield effects associated with QTL for canopy development were investigated using data from four rainfed field trials. Four QTL associated with slower canopy closure, improved yield in specific environments, and notably, were not associated with a yield penalty in any environment. This was likely due to optimised timing of water-use and pleiotropic effects on yield component traits, including spike number and spike length. Overall, this study suggests that slower canopy closure is beneficial for durum wheat production in rainfed environments. Selection for traits or loci associated with canopy development may improve yield stability of durum wheat in water limited environments.

## INTRODUCTION

Durum wheat, or pasta wheat (*Triticum turgidum* L. ssp. *Durum*; 2n = 4x = 28) is an ancient food crop and an important industry in Mediterranean and sub-tropical agricultural regions (Sall et al., 2019). Production is often constrained by drought, during and post anthesis (Loss & Siddique, 1994). This can greatly reduce yield potential by limiting grain number and weight (Gevrek & Atasoy, 2012; Royo, Abaza, Blanco, & del Moral, 2000). Therefore, traits with phenotypic plasticity are important for increasing crop productivity, as trait plasticity allows for adaptive potential in response to environmental variations (Borrell et al., 2014; Maccaferri et al., 2008; Matesanz et al., 2020; Shavrukov et al., 2017). Traditionally, durum wheat breeders have selected for earlier flowering (Bassi & Nachit, 2019; De Vita et al., 2007; Miralles, Rharrabti, Royo, Villegas, & Garcıa del Moral, 2002; Motzo, Giunta, & Pruneddu, 2010) to minimise the impact of end-of-season drought on reproduction and grain-filling.

In addition to optimising flowering time, canopy traits associated with improved water use efficiency can be targeted. For instance, changes in canopy development (e.g., reduction in leaf size or tillering) provide an advantage under terminal drought conditions by shifting water use from pre-to post-anthesis (Borrell et al., 2014; George-Jaeggli, Mortlock, & Borrell, 2017). Limited transpiration rate at high evaporative demand can also conserve water for critical stages later in crop development (Collins, Chapman, Hammer, & Chenu, 2021). There is a fine balance between water supply and demand in crops and as such, the timing of water availability must be matched with phenological development. Although rapid canopy development can increase light interception (Regan, Siddique, Tennant, & Abrecht, 1997) and reduce soil evaporation (Lopez-Castaneda, Richards, & Farquhar, 1995), if there is insufficient stored soil moisture or in-crop rainfall, excessive canopy size may prematurely deplete soil water and exacerbate terminal drought (Nuttall et al., 2012). Thus, crop performance under drought conditions depends on complex source-sink dynamics between carbohydrate and water balance, where there are trade-offs between stress resilience and yield (Collins et al., 2021; Rodrigues, Inze, Nelissen, & Saibo, 2019). Given the dynamic nature of the environment, understanding canopy development may help to identify integrative traits that support yield.

Normalized difference vegetation index (NDVI) is used to characterise canopy attributes and is considered a good surrogate for biomass accumulation, canopy cover and plant vigour (Cabrera-Bosquet et al., 2011; Carlson & Ripley, 1997; Mullan & Reynolds, 2010; Xue & Su, 2017). NDVI, computed as the difference between near-infrared reflectance and red absorption divided by their sum, is influenced by leaf chlorophyll content and canopy architecture (Gamon et al., 1995). NDVI can be measured in a non-subjective and efficient manner which facilitates its use at the field level. The generalized NDVI profile captured during the growing season includes: (1) the green-up phase before canopy closure, also known as the exponential phase; (2) the peak canopy cover phase; and (3) the decline phase as leaves senesce (Brown & de Beurs, 2008; Masialeti, Egbert, & Wardlow, 2010; Smith, Adams, Stephens, & Hick, 1995; Soltani & Galeshi, 2002). Hereafter, the term canopy development refers to the green-up and maximum cover phases.

While many studies have used NDVI to characterize canopy dynamics during the senescence phase (Christopher, Christopher, Borrell, Fletcher, & Chenu, 2016; Christopher et al., 2014; Lopes & Reynolds, 2012; Pinto, Lopes, Collins, & Reynolds, 2016), few have explored canopy development and assessed its impact on yield. In durum wheat, genetic studies have mapped quantitative trait loci (QTL) using NDVI captured at certain developmental stages or specific time-points (Condorelli et al., 2018; Shi et al., 2017). However, NDVI captured at a specific time point does not account for the temporal dynamics of canopy development. This is important to consider, as the correlation between NDVI captured at a specific time-point and yield is strongly dependent on the growth stage (Goodwin, Lindsey, Harrison, & Paul, 2018; Smith et al., 1995; Teal et al., 2006).

Alternatively, NDVI time-series data can be modelled, and features of the growth curve used to study the underlying genetics. In bread wheat and maize, longitudinal growth data has successfully captured trait development over time to reinforce QTL mapping power (Kwak, Moore, Spalding, & Broman, 2016; Lyra et al., 2020; Miao, Xu, Liu, Schnable, & Schnable, 2020; Muraya et al., 2017). Different parameters of the growth curve related to the time period of interest, may be used to describe temporal NDVI dynamics, such as curve threshold values, inflection points and integrals (Bustos-Korts et al., 2019; Christopher et al., 2014; Lopes & Reynolds, 2012; Pinto et al., 2016). Considering the many environmental factors that can affect the canopy status, area under the respective curve summarises the cumulative changes and can provide a general assessment of canopy development. The approach is yet to be applied to study canopy development in durum wheat.

Understanding the genetics of adaptive traits like canopy development, and their interaction with the environment, is critical to support the development of new cultivars with improved adaptation (Hammer et al., 2020). Using a nested-association mapping population, we reveal the genetic components of canopy development in durum wheat. To gain biological and physiological insights into NDVI time-sequential data, we first explored relationships between NDVI, phenology, canopy cover and features of the growth curve. Secondly, we performed association mapping using both time-point NDVI and the area under the curve (AUC) for NDVI. Finally, markers associated with canopy development were used in a linear mixed model approach to investigate marker × environment interactions and yield effects across multiple rainfed environments in Australia.

## MATERIALS AND METHODS

### Plant materials and genotyping

This study examined subsets of a durum wheat nested-association mapping (NAM) population developed at The University of Queensland, as described by Alahmad et al. (2019). The NAM population comprised 920 lines (10 families) generated by crossing eight elite lines from ICARDA Morocco (i.e. Fastoz2, Fastoz3, Fastoz6, Fastoz7, Fastoz8, Fastoz10, Outrob4 and Fadda98) as ‘founders’ to the Australian durum wheat cultivars Jandaroi and DBA Aurora, which served as ‘reference’ varieties. The founder lines were used as donors for drought adaptive attributes in durum wheat breeding programs in the Middle East and North Africa. The reference varieties are preferred by the pasta industry for their quality and therefore widely grown in Australia. The NAM population was genotyped using Diversity Arrays Technology genotyping-by-sequencing single nucleotide polymorphism (SNP) markers (Alahmad et al., 2019). Allele coding used 0, 1, and 2, where 0 is the reference allele homozygote, 1 is the SNP allele homozygote and 2 is the heterozygote.

### Field trials

A subset of the durum wheat NAM population and a selection of Australian durum wheat varieties were evaluated in four rainfed field trials conducted in Australia between 2017 and 2020 (Supplementary Table S1, Table 1), namely “2017_RW” at Roseworthy (34.30 °S; 138.41 °E), South Australia; “2019_TS” at Tosari, near Tummaville (27.51 °S; 151.27 °E), Queensland; and “2017_WW” and “2020_WW” at Warwick (28. 12 °S; 152.06 °E), Queensland. The trials adopted partially replicated row-column designs (Cullis, Smith, & Coombes, 2006), with the exception of 2017_RW which used a randomized complete block design with three replicates. The total number of genotypes ranged from 147 to 309 across the trials, with pairs of trials having between 51 and 146 genotypes in common. For all trials, starting fertilizer was applied at sowing so that nutrients were not limited, and weeds and insects were controlled as required. Based on the nearest weather station to each trial site, weather information was acquired from the Bureau of Meteorology (http://www.bom.gov.au/) and the SILO database (Jeffrey, Carter, Moodie, & Beswick, 2001).

**Table 1.** Attributes of the four rainfed yield trials conducted between 2017 and 2020.

| **Trial** | **2017_RW** | **2017_WW** | **2019_TS** | **2020_WW** |
| --- | --- | --- | --- | --- |
| Location | Roseworthy | Warwick | Tosari | Warwick |
| SD | 9/05/2017 | 22/06/2017 | 8/07/2019 | 1/07/2020 |
| HarvD | 22/11/2017 | 15/11/2017 | 15/11/2019 | 26/11/2020 |
| No. of plots | 576 | 336 | 440 | 456 |
| ICR (mm) | 320.7 | 137.5 | 12.8 | 185 |
| GDD (°C) | 2468 | 2190 | 2222 | 2377 |
| Range of DTF | na | 96-108 | 84-102 | 90-104 |
| Yield (t ha<sup>-1</sup>) | 3.95 | 4.91 | 2.24 | 3.87 |
| No. of genotype | 149 | 147 | 217 | 309 |
| Traits measured | Yield | SL, SN,<br>DTF, Yield | DTF, Yield | Canopy related traits <sup>a</sup>, SL,<br>SN, DTF, PH, Yield |

<sup>a</sup> Canopy related traits include normalized difference vegetation index (NDVI), and fractional green canopy cover (FGCC) at multiple time points at population level, and tiller number measured on 11 selected genotypes. SD, sowing date; HarvD, harvest date; ICR, in-crop rainfall; GDD, growing degree days during growing period; DTF, days to flowering; SL, spike length measured in cm; SN, spike number per square meter; PH, plant height measured in cm. na, data is not available.

To monitor soil moisture and estimate the impact of drought stress on yield in the 2020_WW trial, two check genotypes (DBA Aurora and Fadda98) were sown in dryland and irrigated blocks next to the main experiment. Both blocks were sown on the same day as the main trial. The rainfed block was adjacent to 2020_WW and the irrigation block was adjacent to the rainfed block (separated by buffer rows to prevent lateral movement of soil moisture across treatments). Plot size was consistent with the main trial and each treatment was sown in a completely randomized block design, using 12 replicates in the dryland treatment and 6 replicates in the irrigation treatment. About 20 mm of water was applied through drip tape irrigation every 1-2 weeks to ensure a stress-free growing environment. To determine soil water availability, soil water content was measured for both dryland and irrigated blocks at one week pre-anthesis and at anthesis. In each strip, two soil cores were collected from DBA Aurora and Fadda98 plots in both treatments. Each core was divided into 20cm soil layers: 0–20, 20–40, 40–60, 60–80, 80–100, and 100–120 cm. A subsample from each soil layer was immediately weighed to obtain fresh weight, and then dried to a constant weight at 105 °C. Soil water content for each layer was calculated as [(fresh weight-dry weight)/dry weight]×100%.

### Data collection

The 2020_WW trial was subjected to intensive canopy phenotyping and resulting phenotypes were used for association mapping. The trial comprised 309 genotypes evaluated using a p-rep design (~38%) of 456 plots (6 m × 1.05 m, 4 rows) (Table 2). NDVI data was captured for each plot every 1-2 weeks, specifically 22, 29, 36, 43, 50, 63, 70, 78, 85, 91, 99 and 106 days after sowing (DAS) using a GreenSeeker™ handheld sensor (NTech Industries, Ukiah, CA, USA). Measurements were recorded on sunny and still days, by holding the sensor at approximately 0.6 m height above the crop canopy of the central two rows while walking through the crop at a constant rate. Canopy cover images were also captured for all plots using a mobile phone camera (Apple iPhone10), at 29, 36, 43 and 50 DAS. The RGB images were processed using Canopeo in the Matlab environment, for calculating fractional green canopy cover (FGCC) that measures the canopy surface area (Patrignani & Ochsner, 2015). In each plot, the number of spikes was manually recorded for an inner row (1 m length) to determine spike number per square meter (SN) and spike length (SL) was recorded for six plants. Plant height (PH) of three random plants in each plot was measured at maturity from ground to the top of the spike, excluding the awn length. Flowering time (DTF) was recorded as DAS to 50% flowering (Zadok’s growth stage 65) of all plants in a plot (Zadoks, Chang, & Konzak, 1974). The crop was harvested using a small-plot machine harvester to obtain yield data.

**Table 2.**
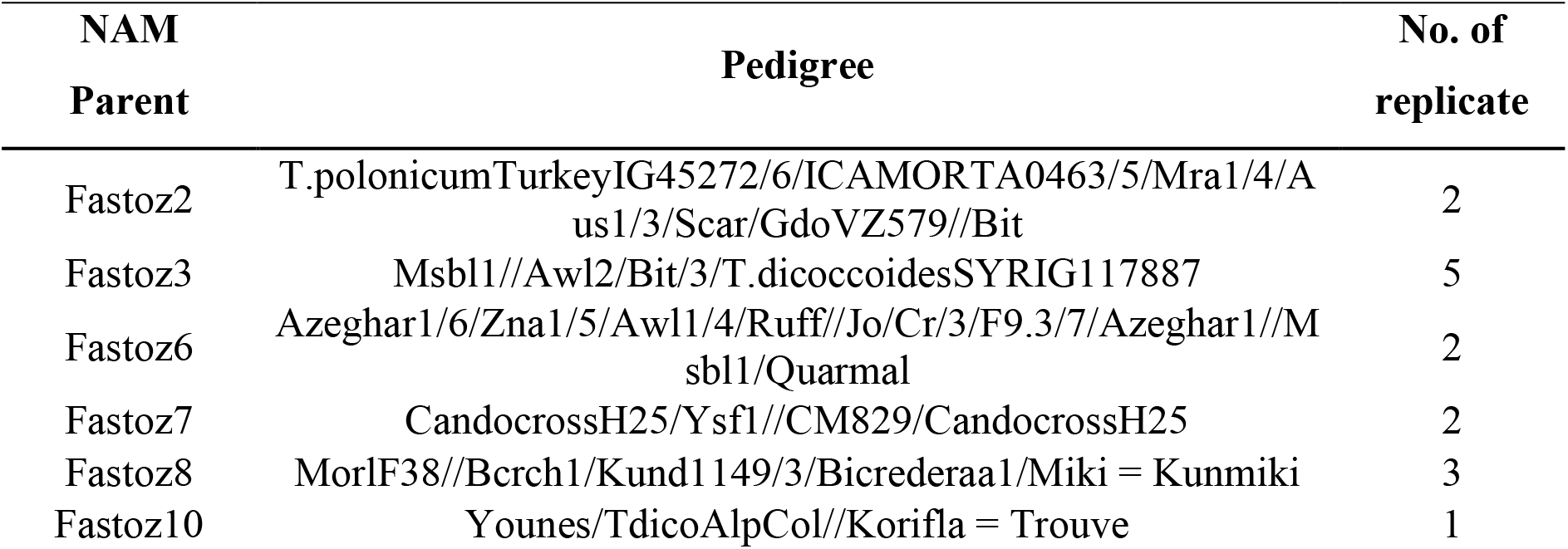
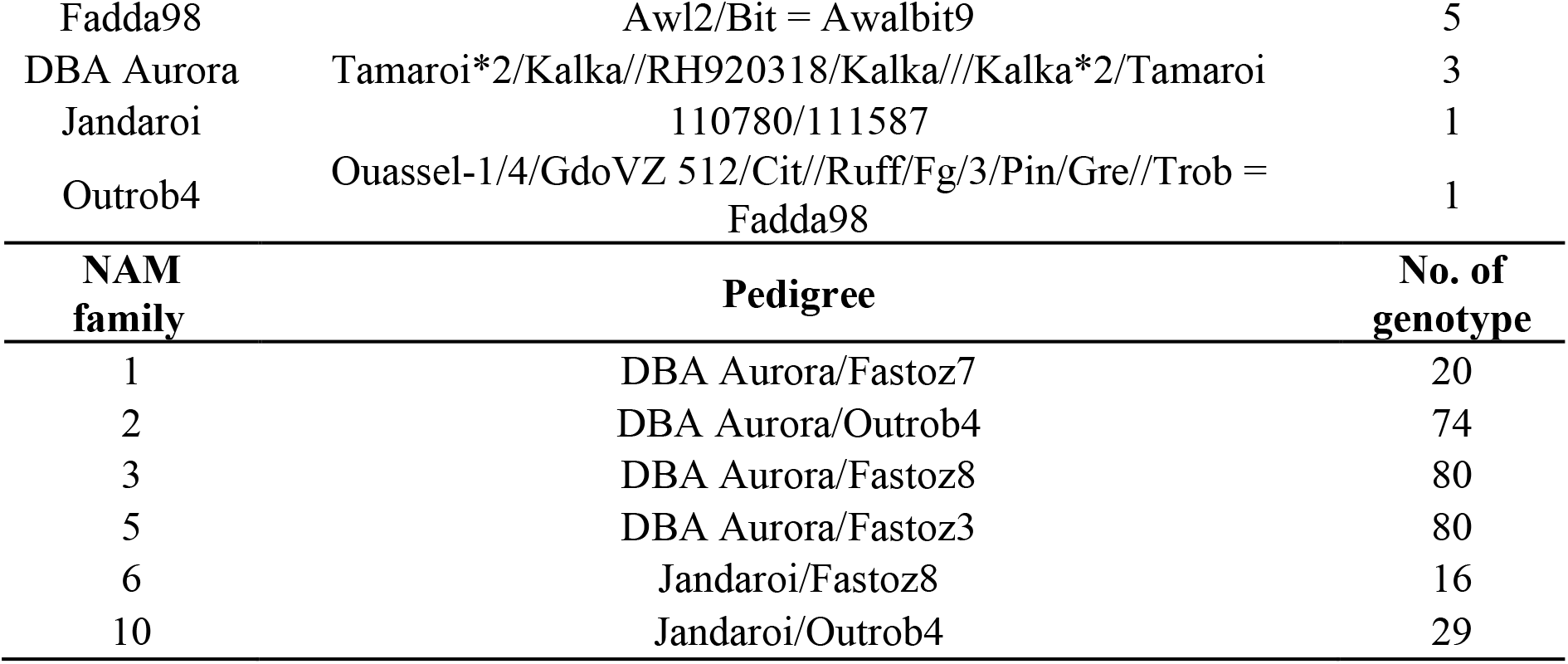
Description of durum wheat genotypes in the 2020 field trial, including the 10 parents of nested-association mapping (NAM) population and subset of 299 NAM lines.

To investigate the relationship between NDVI and crop developmental stages in the 2020_WW trial, a small panel of lines were selected for growth stage tracking from sowing to flag leaf emergence. The panel comprised 11 genotypes, which were selected based on divergent yield performance in previous rainfed yield trials (i.e., high yielding and low yielding lines). Each genotype was replicated 2-3 times in the trial (total 23 plots). Each plot was monitored for Zadoks’ growth stages (GS) and 10 plants in the middle two rows of each plot were tagged for tracking tiller number until flag leaf emergence at 16, 22, 29, 36, 43, 50, 57, 63, 70, 78, and 85 DAS.

To investigate yield effects of SNPs associated with canopy development, analyses used yield data from the 2020_WW trial, plus data from the three other yield (2017_WW, 2017_RW and 2019_TS; Table 1). DTF was captured in all trials, except for 2017_RW (Table 1).

### Analysis of phenotypic data

Spatial analyses were conducted for each trait to correct for spatial heterogeneity within each trial. A linear mixed model was fitted in ASReml-R to estimate adjusted genotype means (best linear unbiased estimates; BLUEs) for all traits in each trial as follows (Butler, Cullis, Gilmour, & Gogel, 2009):

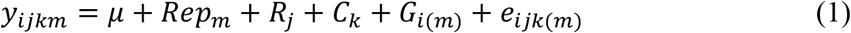

where *y_ijkm_* denotes the plot observation of genotype *i* in replicate *m*, row *j* and column *k*, was modelled by fitting fixed effects for the overall mean (*μ*) and genotype *i* (*G*_*i*(*m*)_); and random effects for replicate *m* (*Rep*_*m*_), row *j* (*R_j_*) and column *k* (*C_k_*); and *e*_*ijk*(m)_ represents the vector of spatially correlated residuals. The variance components of *R_j_*, *C_k_* and *e*_*ijk*(m)_ were assumed to follow 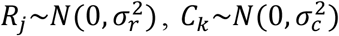 and *e*_*ijk*(m)_ ~ N(0,AR1 ⊗ AR1 σ^2^), respectively. To correct for known or expected sources of variation that were suspected to have some effects on traits, the model was tested to assess the need for fitting of covariates (i.e., differences in establishment between genotypes, lodging). The covariates and random terms were evaluated with Wald chi-squared test and likelihood ratio test, respectively. The model was adjusted according to the identified significant terms at α = 0.05. Except for the replicate, non-significant model terms were dropped in an attempt to obtain the best fit. Slight modifications were made for analysing 2017_RW, where the residual term was modelled by a two-dimensional spatial model with correlation in row direction only.

Time-series modelling of canopy development used NDVI recorded from sowing to the peak of NDVI measures at 78 DAS (Figure 1). To describe the trend of longitudinal BLUEs for NDVI, a smoothing spline was implemented in ASreml-R (Verbyla, Cullis, Kenward, & Welham, 1999). To summarize the NDVI growth curve, AUC was calculated using the following formula:

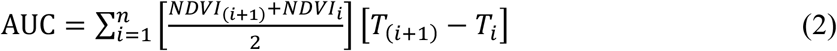

Where *NDVL_i_* is the NDVI prediction at the *i*th DAS; *T*_*i*_ is the *i*^th^ DAS; and *n* is the number of DAS of interest after i.

**Figure 1.**
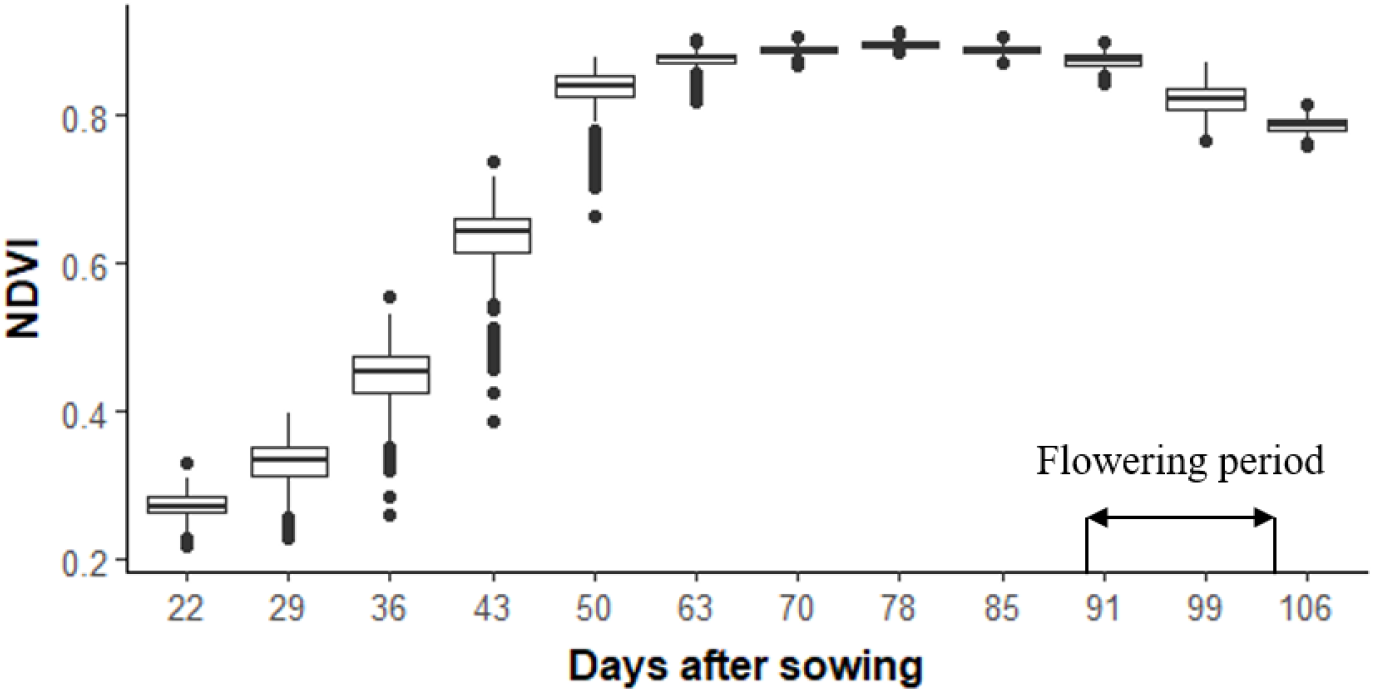
Variability of normalized difference vegetation index (NDVI) of the durum nested-association mapping population in 2020_WW. Each boxplot represents the range of best linear unbiased estimates (BLUEs) for NDVI at each time-point.

### Association mapping

All data captured at the 2020_WW trial were used to perform association mapping. Genotype data was subjected to quality control, which excluded genotypes with > 20% missing marker information and markers with a call rate < 90% and a minor allele frequency (MAF) < 5%. After filtering, a total of 5,298 markers remained for subsequent analyses.

Population structure was investigated using principal component analysis (PCA). An appropriate number of principal components (PCs) were determined by estimating the variances of PC scores. The retained PCs were included as covariates in association tests carried out with “FarmCPU” in R (Liu, Huang, Fan, Buckler, & Zhang, 2016). The *p*‐values of marker–trait associations (MTA) were adjusted in a multiple comparison procedure using false discovery rate (FDR) (Benjamini & Hochberg, 1995). Only associations with adjusted *p*‐values (*p*‐FDR) less than 0.05 were considered as statistically significant and reported.

Analysis of phenotype data revealed correlations between canopy development traits and DTF. Thus, GWAS was initially performed for DTF, PH, spike traits and grain yield. SNPs associated with DTF were then removed from the dataset to ensure that subsequent GWAS performed for canopy developmental traits were able to identify loci independent of flowering behaviour. Two different approaches were used to map QTL for canopy development, the first used BLUEs for NDVI at each time-point whereas the second used AUC derived from spline modelling of time series NDVI.

For each QTL, the positive allele was determined as the allele associated with an increase in trait value. Data from each homozygous allele were tested for normality and homogeneity of variance. The means of genotypes carrying different homozygous alleles were statistically compared by independent t-tests. In several cases where data did not meet the normality criteria, non-parametric Wilcoxon rank sum test was performed to compare the allelic effect on traits.

### Marker × environment analysis

The marker × environment interaction (M × E) was analysed with a linear mixed model, in a multi-environment context using all four field trials (Table 1):

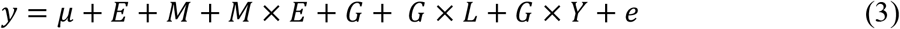

where *y* is the vector of yield BLUEs; *μ* is the general mean; *E* represents trial; *M* denotes SNP; *M* × *E* is the interaction term between SNP and trial; *G* is genotype; *G* × *L* is the genotype by location interaction; *G* × *Y* is the genotype by year interaction, and *e* is the residual, assumed independent with identical distribution. In the model, *E*, *M*, and *M* × *E* were fixed effects, whereas *G*, *G* × *L* and *G* × *Y* were treated as random effects.

The SNP effect was modelled as the sum of a main effect common to all tested environments (M), plus the interaction term representing environments-specific deviations (M × E). Since M × E was tested conditional on the main effect, no attempt was made to interpret the SNP main effect when M × E is significant (Malosetti, Ribaut, & van Eeuwijk, 2013). When M × E is not significant, the SNP main effect could be sufficient to represent the SNP effect. After testing, only SNPs with either significant main effect or M × E effects were reported and further investigated. The predicted means of each SNP allele × trial combination were compared with Tukey’s HSD test.

A summary of the key steps and workflow involved from modelling NDVI time series data to the M × E analysis is provided in Figure 2.

**Figure 2.**
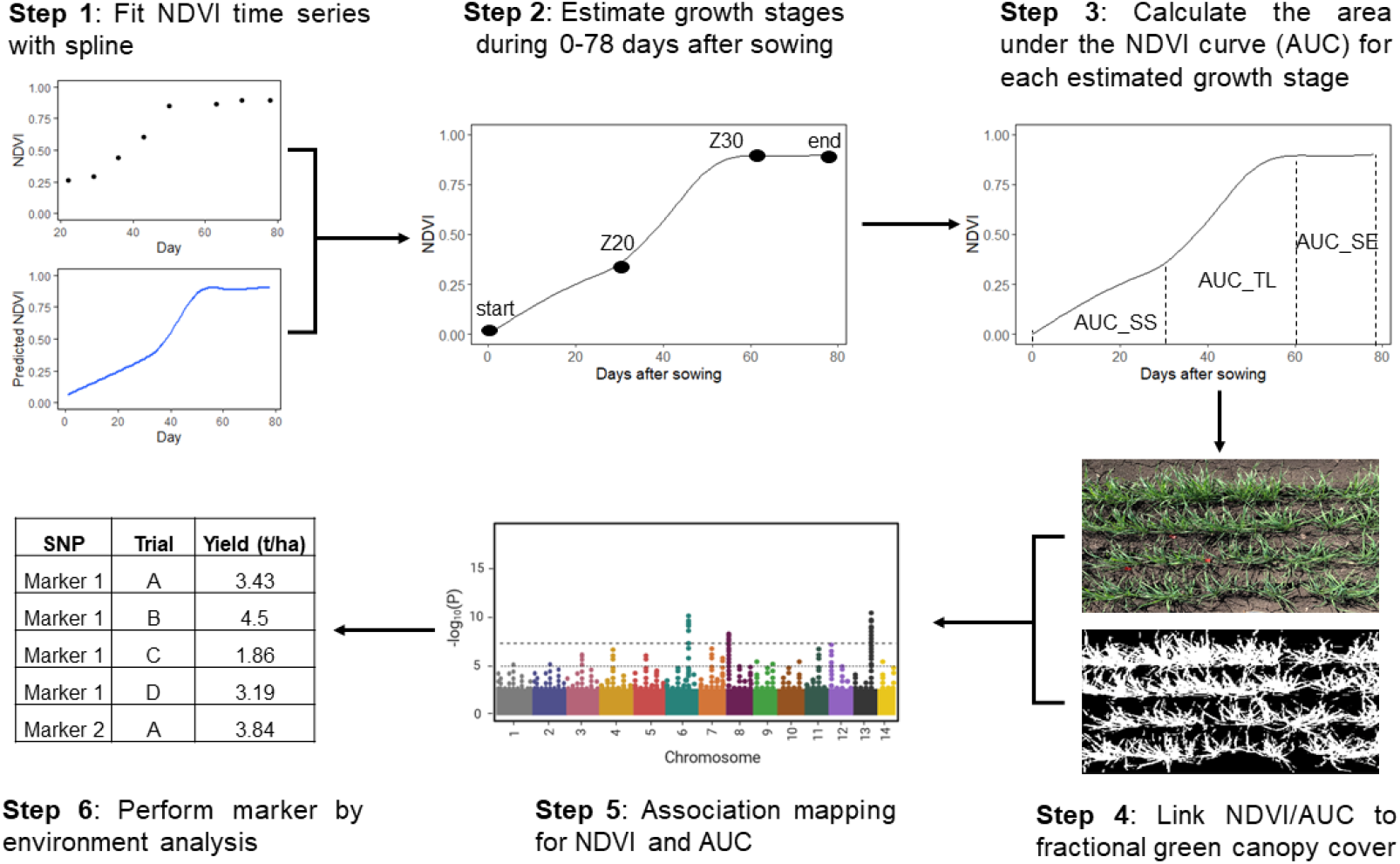
Experimental analyses performed in this study involved the fitting of NDVI time series with a spline (1), estimation of growth stages (2), calculation of AUC for NDVI (3), link of NDVI/AUC to fractional green canopy cover (4), association mapping (5) and marker by environment analysis (6).

## RESULTS

### Field environments experienced variable rainfall patterns

The amount of in-crop rainfall varied across the four trials, ranging from 12.8 mm (2019_TS) to 320.7 mm (2017_RW). While 2019_TS received the least in-crop rainfall (only 12.8 mm) the trial was sown into a full soil profile and the soil is described as deep with high water-holding capacity. This delayed water stress until the grain filling period and the site recorded the lowest mean yield (2.24 t ha^-1^) in comparison with other trials (Supplemental Figure S1C, Table 1). The distribution of rainfall through the season also varied across the trials (Supplemental Figure S1). For example, 2017_RW experienced a typical Mediterranean-type environment, where most rainfall occurred early in the season. In contrast, the two trials conducted at Warwick received more rainfall during the critical grain filling period (Supplemental Figure S1B, D). The highest average yield was obtained in 2017_WW, which was on average 1 tonne ha^-1^ higher than 2017_RW and 2020_WW, and 2.7 t ha^-1^ higher than 2019_TS (Table 1).

To quantify the degree of water stress in the 2020_WW canopy phenotyping trial, soil cores were sampled from dryland and irrigated strips adjacent to the main trial. Sampling performed one week prior to anthesis revealed significant differences in soil moisture (all soil depths from 0 - 1.2 m) for the dryland treatment compared to the irrigated treatment (Supplemental Figure S2A). At anthesis, soil moisture was further depleted, particularly at depth (Supplemental Figure S2A). DBA Aurora and Fadda98 obtained significantly higher yield in the irrigated treatment (Supplemental Figure S2B). Based on average yield differences between dryland and irrigated treatments, the degree of water stress experienced in the 2020_WW trial resulted in an approximate average yield loss of 1.1 t ha^-1^.

### Relationships between NDVI, phenology traits and yield-related traits

The 309 NAM genotypes evaluated in 2020_WW trial displayed a high degree of phenotypic variation for temporal NDVI (Figure 1). Phenotypic variation in NDVI was largest at the beginning of the growing season, reached a peak at 36 DAS and reduced thereafter. The NDVI reached an average peak value of 0.9 at 78 DAS (Table 3). The Pearson correlation among all traits was computed (Figure 3). For a specific time-point NDVI, its correlations with other timepoints decreased with increasing developmental stage. Positive correlations between NDVI and SL were significant at 29 and 78 DAS (*p* < 0.05). A reverse trend was observed for SN, where NDVI in the early season (22 and 29 DAS) was negatively correlated with SN. For plant height, the only correlation with NDVI was evident at 50 DAS (*p* < 0.05). Further, NDVI captured over time was negatively correlated with DTF (Figure 3), with higher NDVI associated with faster time to flowering. No direct NDVI-yield relationship was found before 63 DAS in our study. Rather, NDVI recorded closer to flowering time was more highly related to final yield, as the strongest correlation between NDVI and yield was observed at 78 DAS (*p* < 0.001).

**Table 3.** Summary statistics for traits studied in the 2020 field trial, including minimum (Min), mean, maximum (Max), and coefficient of variation (CV) for trait best linear unbiased estimates (BLUE).

| Traits <sup>a</sup> | Min | Mean | Max | CV (%) |
| --- | --- | --- | --- | --- |
| NDVI_22DAS | 0.22 | 0.27 | 0.33 | 6.3 |
| NDVI_29DAS | 0.23 | 0.33 | 0.4 | 9.3 |
| NDVI_36DAS | 0.26 | 0.45 | 0.55 | 9.6 |
| NDVI_43DAS | 0.39 | 0.63 | 0.74 | 7.9 |
| NDVI_50DAS | 0.67 | 0.84 | 0.88 | 3.5 |
| NDVI_63DAS | 0.82 | 0.88 | 0.9 | 1.3 |
| NDVI_70DAS | 0.87 | 0.89 | 0.91 | 0.6 |
| NDVI_78DAS | 0.88 | 0.9 | 0.91 | 0.5 |
| FGCC_29DAS | 3.21 | 10.35 | 16.52 | 17.1 |
| FGCC_36DAS | 7.6 | 23.34 | 33.67 | 16.9 |
| FGCC_43DAS | 20.63 | 44.01 | 57.51 | 13.5 |
| FGCC_50DAS | 39.81 | 70.86 | 83.91 | 9.9 |
| AUC_SS | 3.77 | 5.6 | 7.08 | 7.4 |
| AUC_TL | 14.83 | 20 | 21.8 | 5 |
| AUC_SE | 15.47 | 15.98 | 16.32 | 0.7 |
| AUC_VS | 34.99 | 41.57 | 44.46 | 3.4 |
| PH (cm) | 58 | 77 | 94 | 7.9 |
| DTF (days) | 90 | 95 | 104 | 2.1 |
| SL (cm) | 7.19 | 9.21 | 11.35 | 8.3 |
| SN (-m<sup>-2</sup>) | 115 | 248 | 365 | 17.7 |
| Yield (t-ha<sup>-1</sup>) | 1.66 | 3.63 | 5.56 | 17.8 |

<sup>a</sup>DAS, days after sowing; FGCC, fractional green canopy cover; AUC, area under the NDVI curve; SS, seedling stage; TL, tillering; SE, stem elongation; VS, vegetative stage.

**Figure 3.**
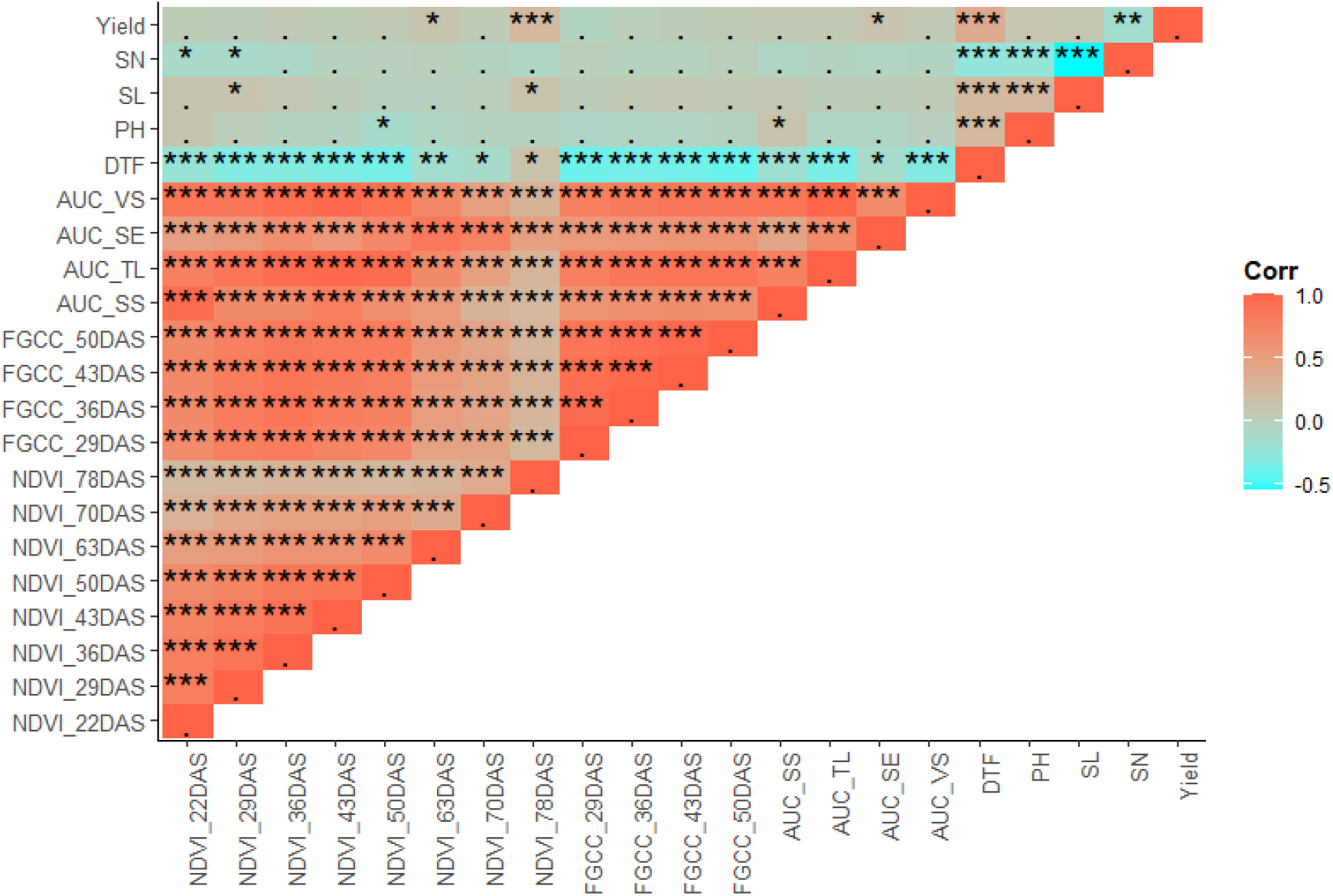
Heatmap of trait by trait correlations in the 2020_WW trial. Pearson’s correlation was computed for each pair of traits. The colour key represents the Pearson’s correlation coefficient. Level of significance *: *p* < 0.05; **: *p* < 0.01; ***: *p* < 0.001. The explanation for trait abbreviation can be found in Table 3.

### Modelling NDVI over time to estimate growth stages

The longitudinal data fitted with a spline showed that the NDVI growth curves overall, increased slowly initially, then rapidly, before reaching a final plateau (Figure 4). This suggested three different growth phases likely involved in canopy development in durum wheat. Although the trends of these curves were more or less in parallel over time, the distribution of genotype-specific NDVI trajectories indicated some heterogeneity in each phase, leading to variation in phase-specific AUC traits. For this reason, the entire simulated AUC during the vegetative stage (AUC_VS) could be divided into three phases, each illustrating a different growth status, to capture phase-specific variation that contributes to the overall canopy development.

**Figure 4.**
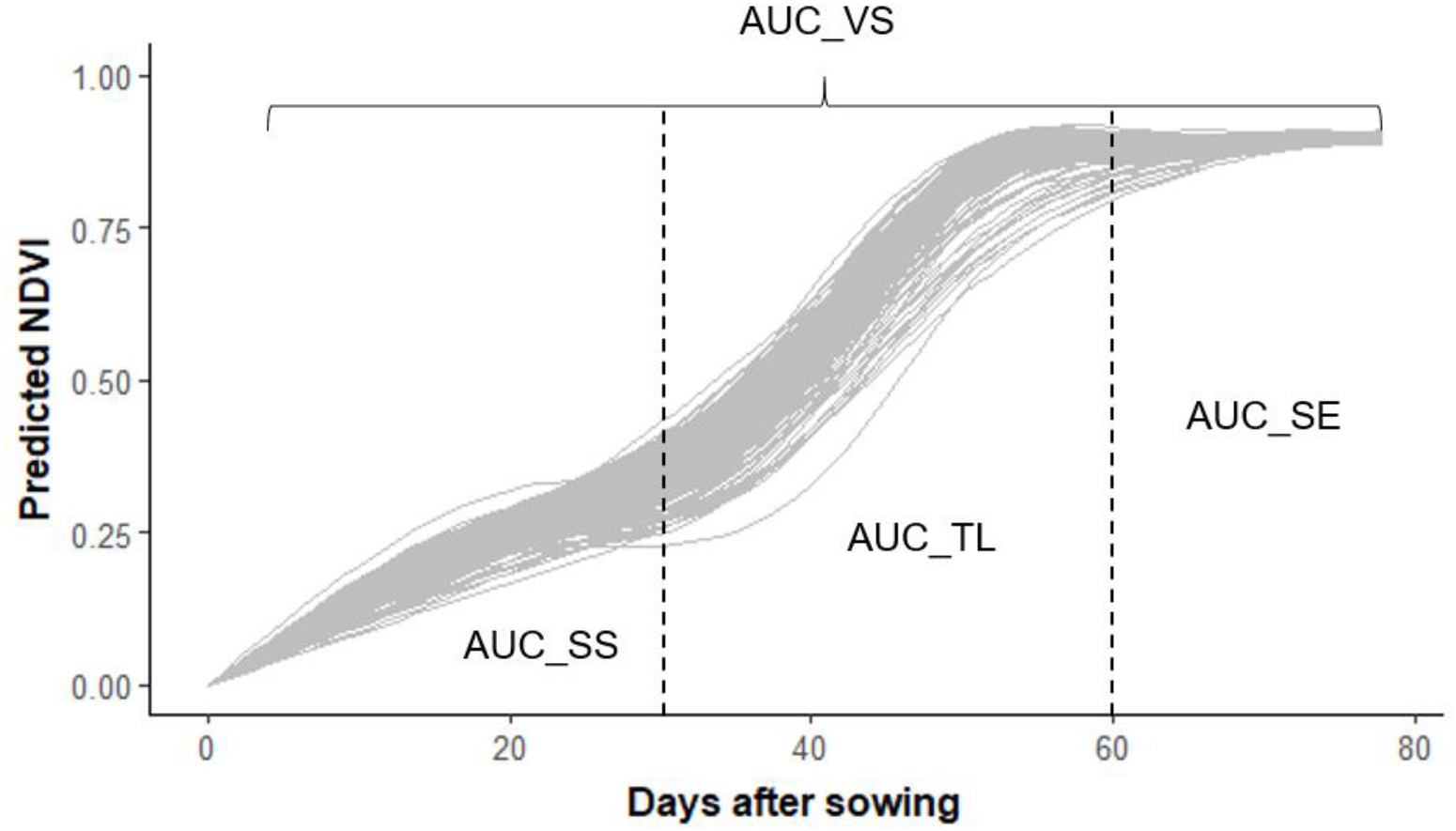
Trajectories of time-series NDVI of all genotypes in the mapping population fitted with smoothing splines. The breakpoints 30 and 60 days after sowing were used to bin the whole range of simulated NDVI data. First phase = seedling stage (AUC_SS, 0-30 DAS), second phase = tillering (AUC_TL, 30-60 DAS), third phase = stem elongation (AUC_SE, 60-78 DAS).

To understand how NDVI dynamics reflected changes in vegetation phenology, 11 genotypes including 10 NAM lines and the reference variety DBA Aurora were investigated. Given the similar phenology of the 11 genotypes, Zadok’s GS20 (start of tillering) and GS31 (first node) were aligned with 29 and 63 DAS (Supplemental Figure S3), respectively. Since no significant change was observed for the tiller number of most genotypes from 57-63 DAS (Supplemental Figure S3), Zadok’s GS30 (start of stem elongation) was estimated at 60 DAS. To evaluate the use of NDVI to define the growth stage, we hereafter used 30 and 60 DAS as two breakpoints to approximate GS20 and GS30, respectively.

Using 30 DAS as the first breakpoint, the NDVI trajectories of genotypes displayed two distinct growth patterns before and after the point. For instance, the sharp increase in NDVI after 30 DAS suggested a transition from seedling to tillering stage. According to the fitted NDVI curves, most genotypes reached the start of maximum canopy cover at approximately 60 DAS (Figure 4). This finding aligned with the start of the maximum canopy cover as indicated by time-point NDVI measures (Figure 1), where NDVI after 63 DAS remained constant. As such, the estimated transition from tillering to stem elongation by the NDVI curve was deemed reasonably accurate. However, to further identify and interpret phenology metrics, the saturation issue may impact NDVI-based recommendations as NDVI becomes insensitive to changes in canopy structure when the crop reaches canopy closure.

### The relationship between time-point NDVI and AUC traits

NDVI curves were binned into three growth stages: seedlin*g* stage (SS, 0-30 DAS), tillering stage (TL, 30-60 DAS) and stem elongation stage (SE, 60-78 DAS) (Figure 4). Accordingly, the AUC traits for each stage were designated AUC_SS, AUC_TL and AUC_SE, and were used to quantify the cumulative status for each stage. Given this, the same duration of each growth stage was applied to all studied genotypes. Hence, differences in growth rate appeared to contribute to variation in AUC, where higher AUC values represented faster canopy development and closure.

As expected, because of the linear nature of the operations involved, stage-specific AUC traits showed strong correlations with NDVI measured within the respective stage (Figure 3). Moreover, stage-specific AUC traits were also found to correlate well with NDVI measured at other stages. The integral NDVI approach ensured that canopy differences related to yield formation were captured. For example, AUC_SE was correlated with yield, but only some of the NDVI time-points during SE showed significant correlations with yield (e.g., 70 DAS was not correlated, but readings captured at 63 DAS and 78 DAS were, as shown in Figure 3). These results highlighted the robustness and suitability of the approach for proceeding with genetic dissection studies.

### Time-point NDVI and AUC correlate with canopy cover

NDVI displayed a positive linear relationship with FGCC before NDVI reached the maximum value of 0.9 (Figure 5). Most genotypes obtained 80% FGCC at 50 DAS. Thus, rapid growth during the tillering stage could almost achieve canopy closure before the start of stem elongation. Moreover, all AUC traits showed significant correlations with FGCC, except for AUC_SS (Figure 3). This highlights the value of NDVI to estimate canopy cover as measures were similar to RGB-based estimates. As such, a higher NDVI and/or a greater AUC value represented a larger canopy that was faster to close.

**Figure 5.**
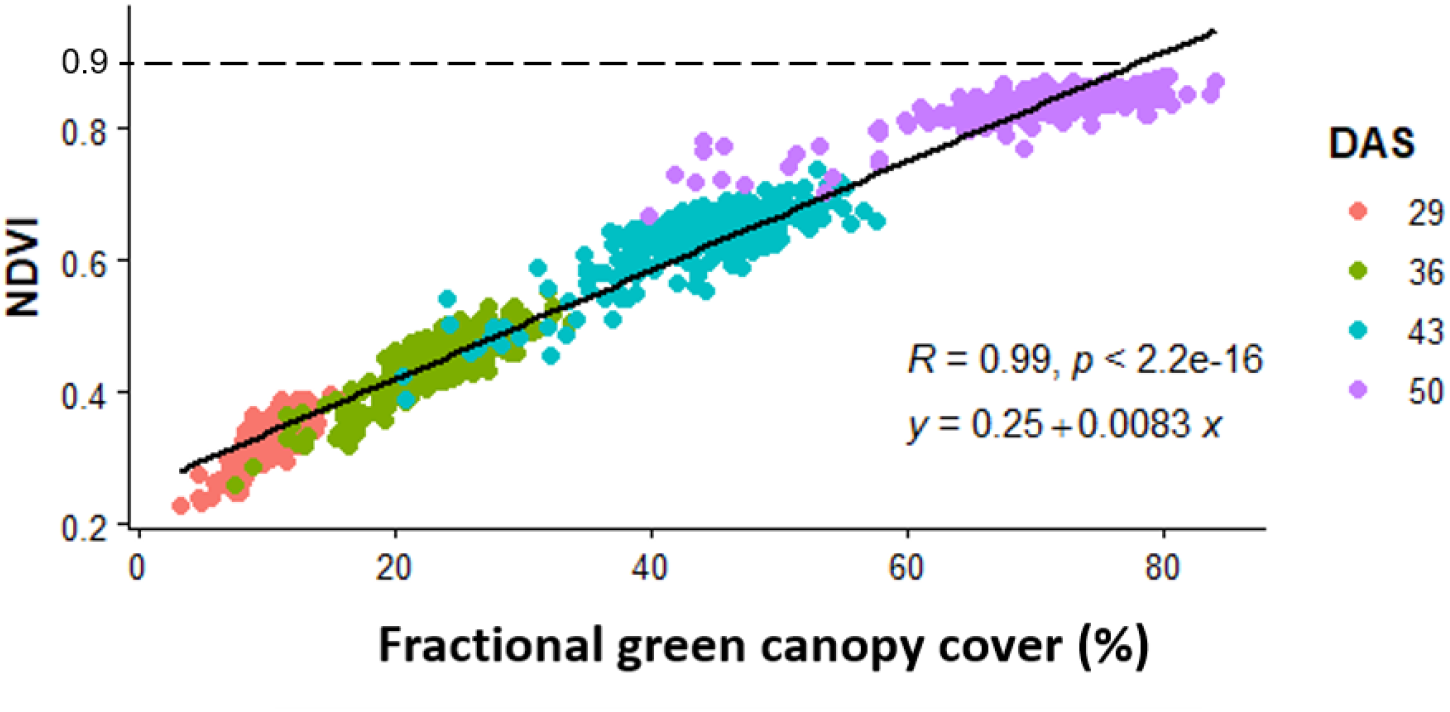
Relationship between BLUEs of NDVI and fractional green canopy cover measured at 29, 36, 43 and 50 DAS.

### Association mapping for canopy development

The PCA revealed six clusters in the NAM population (Supplemental Figure S4A), which aligned with the family structure (Table 2). The first five PCs were used as covariates in association mapping, because explained variance rapidly decreased until PC = 5 and changed little thereafter (Supplemental Figure S4B). The first two PCs explained ~ 23% of the genetic variance (Supplemental Figure S4B).

GWAS for DTF using 5,298 high quality SNPs identified a total of 33 SNPs associated with flowering time and 6 SNPs associated with PH (Supplementary Table S2). A total of 7 SNPs across 5 chromosomes were associated with spike length (Supplementary Table S2). Following removal of the 33 SNPs associated with flowering time, GWAS was performed using a subset of 5,265 SNP markers for time-point NDVI and stage-based AUC captured in the 2020_WW trial. Using time-point NDVI, a total of 7 significant MTAs were detected across nine chromosomes, including 2A, 2B, 4A, 6A and 7A (Table 4). Among these, only one SNP was detected for more than one NDVI time-point (i.e., SNP 1271404 on chromosome 2A). Notably, in agreement with the genetic variation in NDVI (Figure 1), most MTAs were identified at specific time-points between 29 and 50 DAS.

**Table 4.** Summary of results from association mapping of canopy development traits in the 2020 field trial (2020_WW).

| Trait | SNP | Positive allele | Chr | MAF | Pos.St (bp) | Pos.End (bp) | -log<sub>10</sub> (*p*) | -log<sub>10</sub>(*p*-FDR) |
| --- | --- | --- | --- | --- | --- | --- | --- | --- |
| NDVI_29DAS | 1271404 | 1 | 2A | 0.21 | 745108704 | 745108769 | 6.51 | 2.79 |
|  | 5324123 | 1 | 4A | 0.13 | 642498972 | 642499004 | 4.83 | 1.59 |
|  | 4404447 | 1 | 6A | 0.35 | 12144868 | 12144909 | 5.05 | 1.64 |
| NDVI_36DAS | 1271404 | 1 | 2A | 0.21 | 745108704 | 745108769 | 5.13 | 1.42 |
| NDVI_43DAS | 1271404 | 1 | 2A | 0.21 | 745108704 | 745108769 | 5.01 | 1.60 |
|  | 1145239 | 1 | 7A | 0.23 | 44750677 | 44750726 | 5.32 | 1.60 |
| NDVI_50DAS | 1271404 | 1 | 2A | 0.21 | 745108704 | 745108769 | 6.65 | 2.93 |
|  | 5324050 | 0 | 2A | 0.12 | 769791063 | 769791095 | 4.83 | 1.60 |
|  | 3949783 | 0 | 2B | 0.28 | 697623388 | 697623452 | 5.38 | 1.97 |
| NDVI_70DAS | 996714 | 1 | 7A | 0.17 | 109908919 | 109908968 | 5.71 | 2.00 |
| AUC_TL | 1271404 | 1 | 2A | 0.21 | 745108704 | 745108769 | 5.60 | 1.88 |
| AUC_SE | 4004899 | 1 | 2A | 0.35 | 735922887 | 735922955 | 4.47 | 1.35 |
|  | 1095539 | 0 | 2B | 0.29 | 618076302 | 618076370 | 7.17 | 3.45 |
|  | 3946360 | 1 | 4A | 0.19 | 24099524 | 24099564 | 5.46 | 2.22 |
|  | 1093322 | 0 | 5B | 0.20 | 528696884 | 528696822 | 5.91 | 2.49 |
| AUC_VS | 1271404 | 1 | 2A | 0.21 | 745108704 | 745108769 | 5.98 | 2.27 |
|  | 3934649 | 1 | 7A | 0.21 | 633603150 | 633603198 | 4.89 | 1.48 |

Chr, chromosome; MAF, minor allele frequency; Pos.St, the start of the SNP position in base pair (bp) on the 'Svevo' durum reference genome; Pos.End, the end of the SNP position on the 'Svevo' durum reference genome; -log<sub>10</sub> (*p*), -log<sub>10</sub> of uncorrected *p* value of marker-trait association; -log<sub>10</sub>(*p*-FDR), -log<sub>10</sub> of *p* value adjusted by false discovery rate.

To identify markers associated with AUC, we conducted association mapping using the following traits: AUC_SS, AUC_TL, AUC_SE and AUC_VS. This detected six significant MTAs including, SNP 1271404 on chromosome 2A which was also detected using time-point NDVI measures during the TL growth stage. Mapping AUC enabled the identification of five additional signals, on chromosome 2A (SNP 4004899), 2B (SNP 1095539), 4A (SNP 3946360), 5B (SNP 1093322) and 7A (SNP 3934649) (Table 4).

### M × E analysis revealed markers associated with grain yield

The M × E interaction analysis was conducted to assess the significance and strength of the SNP effects on yield across trials. Analyses focussed on 12 SNPs that were associated with canopy development and segregating in population subsets evaluated across all trials.

The allelic effects on canopy development were first explored using data collected from 2020_WW. For all SNPs, the allele associated with either higher NDVI or larger AUC was defined as the positive allele, which was linked to rapid canopy closure (Table 4). A linear mixed model approach was employed to evaluate effects for each of the 12 SNPs using yield data from four rainfed trials (Table 1). In the current study, no significant marker main effect was detected for yield. Instead, 9 markers showed significant M × E interactions for yield (Table 5). Notably, SNP 1095539, 3949783, 4404447 and 5324123 showed significant yield effects in 2017_WW, 2017_WW, 2017_RW, and 2020_WW, respectively (Table 5). Alleles associated with a significant yield benefit were associated with slow canopy closure (Supplemental Figure S5; Table 4, 5). Therefore, these four SNPs of interest were subjected to further investigation.

**Table 5.**
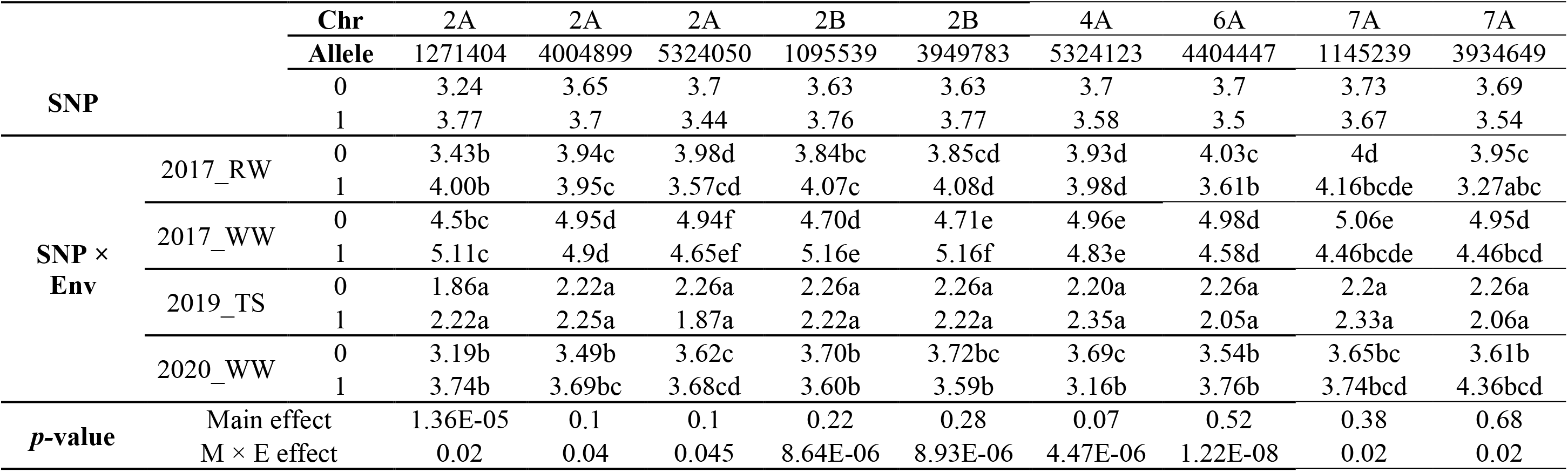
Summary of marker × environment (M × E) interactions for yield. Predicted means of yield are presented for allele 0 and 1 at each SNP locus, and each SNP allele × trial combination. Significant differences are indicated by different letters at 0.01 probability level following Tukey’s test.

### Alleles influencing canopy development and yield were also associated with spike length and spike number

Three of the four marker alleles associated with a slower closing canopy and yield (1095539, 3949783, 4404447 and 5324123) also showed associations with SL or SN. Interestingly, SNP 5324123 was strongly associated with both SL and SN in 2020_WW, but the yield benefit in this trial was related to a reduction in SN (Supplemental Figure S6D, Table 5-6). Similarly, the significant yield effect of SNP 1095539 in 2017_WW was associated with SL (Supplemental Figure S6A, Tables 5-6). On the other hand, SNP 3949783 was associated with SL in 2020_WW but not yield (Tables 5-6). SNP 4404447 was not associated with either component traits (Table 6). Notably, SNP 4404447 was not associated with yield in 2017_WW and 2020_WW and these were the environments where data for SL and SN were captured (Table 5). Overall, alleles associated with slow canopy closure supported yield, however, the contribution and yield benefit associated with pleiotropic effects on SL and SN appeared highly context dependent across the environments.

**Table 6.** Comparison of two homozygous alleles at four SNP loci for spike traits measured in 2017_WW and 2020_WW. Data was analysed with unpaired t-test in two trials separately.

| SNP | Chr | Trial | Allele | N<sup>a</sup> | SL (cm) | *p*-value<br>SL | SN | *p*-value<br>SN |
| --- | --- | --- | --- | --- | --- | --- | --- | --- |
| 1095539 | 2B | 2017_WW | 0 | 80 | 7.29 | <0.05 | 323 | 0.16 |
|  |  |  | 1 | 66 | 7.52 |  | 311 |  |
|  |  | 2020_WW | 0 | 89 | 8.98 | <0.001 | 254 | 0.11 |
|  |  |  | 1 | 218 | 9.3 |  | 245 |  |
| 3949783 | 2B | 2017_WW | 0 | 80 | 7.35 | 0.35 | 321 | 0.40 |
|  |  |  | 1 | 66 | 7.46 |  | 314 |  |
|  |  | 2020_WW | 0 | 84 | 8.99 | <0.001 | 248 | 0.94 |
|  |  |  | 1 | 223 | 9.31 |  | 248 |  |
| 5324123 | 4A | 2017_WW | 0 | 92 | 7.44 | 0.44 | 311 | <0.05 |
|  |  |  | 1 | 50 | 7.34 |  | 330 |  |
|  |  | 2020_WW | 0 | 256 | 9.27 | <0.001 | 245 | <0.01 |
|  |  |  | 1 | 36 | 8.82 |  | 268 |  |
| 4404447 | 6A | 2017_WW | 0 | 122 | 7.39 | 0.37 | 321 | 0.12 |
|  |  |  | 1 | 25 | 7.46 |  | 304 |  |
|  |  | 2020_WW | 0 | 200 | 9.18 | 0.28 | 250 | 0.27 |
|  |  |  | 1 | 109 | 9.28 |  | 244 |  |

<sup>a</sup> Number of genotypes carrying respective SNP allele in the trial.

## DISCUSSION

Wheat yield is determined by the interaction between source, which is the availability of photoassimilates, and sink, which is the number of grains per unit area (Reynolds et al., 2017). Canopy development underpins yield potential, as it influences the capture of light, water use, transpiration, and overall biomass production. In water-limited environments, optimal canopy development balances water use both pre- and post-anthesis to maximise yield. For example, while slow canopy development may not favour biomass accumulation pre-anthesis, it can help to conserve soil moisture for the critical grain filling period. However, in environments where water is non-limiting, optimal canopy development should maximise biomass production and overall sink strength as there is no penalty of high water-use early in the season. This study revealed a high degree of variation for temporal canopy dynamics in elite durum wheat populations derived from Australian × ICARDA crosses, which could be used to improve durum wheat adaptation to a range of target environments. New knowledge of the underlying genetics and value of canopy developmental traits described in this study provide important steps towards the development of new cultivars with improved resource-use efficiency to maximise crop yield.

### Using NDVI to measure canopy development

NDVI serves as an easy-to-measure indicator of canopy development in real time. The link between NDVI and canopy development is strongly underpinned by the functional relationship between NDVI and aboveground biomass. In accordance with previous research, time-specific NDVI measures during the early growing period were generally poor predictors of yield (Magney, Eitel, Huggins, & Vierling, 2016), whereas NDVI captured at the peak of canopy development is more associated with yield. This is somewhat expected due to the complexity underpinning yield development. Furthermore, the stem elongation to flowering phase is considered the most critical for determining grain number and ultimately sink strength.

Phenological and environmental changes over time affect the canopy status represented by NDVI. Therefore, the use of the NDVI integral (i.e., area under the curve) provides an advantage over time specific NDVI as it captures the impact of those changes on canopy development. To accurately assess long-term patterns of canopy development, regular NDVI measurements are required. Nonlinear models have been widely used to account for the complexities of plant growth (Paine et al., 2012; Villegas, Aparicio, Blanco, & Royo, 2001).

Previous studies comparing different models to characterise the dynamics of NDVI over time found that spline-fitting better approximated the variation of smoothed NDVI values than other non-linear functions and was more suitable for describing the time-series model (Sun et al., 2017; Vorobiova & Chernov, 2017). In this study, the trajectories of smoothed NDVI data showed a typical temporal pattern of NDVI evolution during the vegetative stage, where crop emergence was followed by a rapid growth period, then a relatively stable period of maximum vegetation approaching anthesis. Therefore, cumulative NDVI at specific growth stages could be used to gain insights of the physiological drivers underpinning grain yield.

### The genetics of canopy development

To identify loci underpinning canopy development, time-point NDVI data were treated as independent traits and association mapping was performed for each timepoint to identify time-specific NDVI (Figure 2). Next, spline-fitted curve-derived AUC was subject to mapping, and results were compared to mapping of time-point NDVI.

Time-point NDVI measures were highly correlated, suggesting the underlying genetic controls either provide long-term regulation of canopy growth or have prolonged effects originating from an early growth phase. This was further confirmed in the mapping results, where the 2A QTL (SNP 1271404) was detected by multiple time-point NDVI.

AUC in the current study was used to capture the genetic basis of canopy growth with respect to key developmental stages. However, some QTL could only be detected using time-point NDVI and it is unclear why these QTL could not be captured using the AUC approach. One possible explanation is that they were transient QTL sensitive to time of data collection, whereas AUC represented a period of growth, and therefore more likely detected loci with more consistent and robust effects. Thus, time-point NDVI and AUC had different mapping strengths and were complementary to each other. The use of AUC for mapping time-dependent canopy development should be implemented on a case-by-case basis, with the aim of ensuring good quality data for use in modelling canopy dynamics. Additionally, AUC values should not be directly compared across different studies, as AUC is a product of many contributing factors, including the environment, modelling approach, phenotyping method and crop phenology.

This study uncovered four SNPs on chromosomes 2B (1095539 and 3949783), 4A (5324123) and 6A (4404447) that could be useful for durum wheat breeding, as alleles associated with slow canopy closure were linked to a yield advantage in some environments, but not a yield penalty in other environments. Notably, all four SNPs were not associated with DTF, which improves the utility of the genes from a breeding perspective, as canopy development could be manipulated without shifting flowering time. Three of the four SNPs (1095539, 3949783 and 5324123) were also associated with yield component traits: SN and SL, where SL is considered a proxy for the number of grains per spike (Baye, Berihun, Bantayehu, & Derebe, 2020). Interestingly, most of these SNPs were detected using NDVI measures recorded at the early tillering stage, an important phase for spike formation in wheat (Khadka, Earl, Raizada, & Navabi, 2020), when no correlation between NDVI and yield was found. The 2B (3949783) and 6A (4404447) regions have previously been reported to influence a range of spike traits including spike dry matter, grain weight per spike, grains per spike and grain weight (Giunta, De Vita, Mastrangelo, Sanna, & Motzo, 2018; Mangini et al., 2018; Patil et al., 2013; Peleg et al., 2009; Soriano, Malosetti, Rosello, Sorrells, & Royo, 2017). While the 4A region (5324123) has not previously been reported for SN and SL per se, it has been reported to influence similar or related traits, including biomass, harvest index, spike harvest index, spike density (spikelet number/SL), and importantly grain yield (Mengistu et al., 2016; Peleg, Fahima, Korol, Abbo, & Saranga, 2011; Peleg et al., 2009; Tzarfati et al., 2014).

### Yield benefits of slow canopy development, trade-offs and pleiotropic effects

Four QTL for slow canopy development were associated with yield in three of the four rainfed environments (2017_RW, 2017_WW and 2020_WW). These three trials likely experienced water stress at anthesis, whereas 2019_TS received very little in-crop rainfall (Supplemental Figure S1C) and likely experienced pre-anthesis water stress. This highlights the value of slow canopy development in water-limited environments that experience drought at anthesis or during the grain filling period. As discussed above, the benefit of a slow closing canopy likely manifests from water savings that support yield formation during grain filling. Without having sufficient environmental data to perform robust envirotyping in APSIM (Chenu, Deihimfard, & Chapman, 2013), it is difficult to quantify the degree of water-stress in the rainfed experiments. However, soil moisture was measured in the dryland and irrigated strips adjacent to the 2020_WW trial. Soil moisture under the rainfed conditions was clearly depleted at anthesis, particularly in deeper soil layers (Supplemental Figure S2A), and the water limitation resulted in an average yield loss of 1.1 t ha^-1^. This highlights the impact of drought under rainfed conditions in Australia, despite three of the four trials being conducted on deep soils with a high water-holding capacity. Although soil moisture data was not available for 2017_WW, the same site was used for 2020_WW. In 2017_WW the trial received less in-crop rainfall compared to 2020_WW (Table 1), suggesting it likely experienced similar or more severe water stress than 2020_WW. Terminal drought often occurs in Mediterranean-type environments, such as South Australia. In 2017_RW, about 90% of the in-crop rainfall occurred from May to September, suggesting the trial experienced water stress late in the season (Supplemental Figure S1A). Regardless of the delayed onset of water stress (compared with the Warwick sites in Queensland), the four QTL associated with slow canopy development contributed positive yield effects.

Harvested wheat yield is a result of three components: spike number per unit area, grain number per spike, and average grain weight (Simmonds et al., 2014; Zhang et al., 2018). In the current study, the relationship between SL, SN and yield varied across the two trials (2017_WW and 2020_WW). Specifically, SL showed a significant correlation with yield in 2017_WW, but not in 2020_WW (Supplemental Figure S6A, C), while SN showed a significant correlation with yield in 2020_WW, but not in 2017_WW (Supplemental Figure S6B, D). Previous studies in both bread and durum wheat have reported positive relationships between SL and yield under water-limited conditions (Munir, Chowdhry, & Malik, 2007; Nofouzi, 2018). In this scenario, genotypes with fewer tillers could accumulate less biomass, but produce longer spikes with more grains, to achieve a yield advantage.

It is plausible that loci associated with canopy development and yield component traits could be involved in modulating root architecture. The study by Voss-Fels et al. (2018) reported that allelic variation at *Vrn1*, a key gene in the wheat flowering pathway, not only influences spike and canopy development, but also root system architecture. In the current study, a QTL on 2B (1095539) associated with AUC_SE was positioned in close proximity with previously reported QTL for root growth angle and primary root length (Maccaferri et al., 2016). Thus, either closely positioned or pleiotropic loci on chromosome 2B could be important for both above- and below-ground developmental traits.

Clearly, the value of different yield component traits in durum wheat is highly context dependent, and genotypes can exploit a range of pathways to maximise yield in each environment. A priority for future research is to understand the complex interactions and possible pleiotropic effects of loci influencing both canopy development and yield component traits. Such insight will enable selection and deployment of desirable gene combinations in breeding programs seeking to develop new varieties with improved resilience and productivity.

## Core Ideas

- NDVI trajectory traits provided insight into canopy dynamics and could guide durum breeding
- QTL associated with slow canopy closure supported yield in rainfed environments
- Yield benefits associated with canopy QTL were also associated with spike number and length

## Abbreviations

AUC: area under the curve
BLUE: best linear unbiased estimate
DAS: days after sowing
DTF: days to flowering
FDR: false discovery rate
FGCC: fractional green canopy cover
GS: Zadoks’ growth stages
MTA: marker–trait associations
NAM: nested-association mapping
NDVI: normalised difference vegetation index
PC: principal component
PCA: principal component analysis
PH: plant height
QTL: quantitative trait loci
SE: stem elongation
SL: spike length
SN: spike number per square meter
SNP: single nucleotide polymorphism
SS: seedling stage
TL: tillering
VS: vegetative stage

## ACKNOWLEDGEMENT

Funding was supported by Grain Research and Development Corporation (GRDC; project code UOQ1903-007RTX). We are grateful for technical support and field trial maintenance provided by the Queensland Department of Agriculture and Fisheries.

## SUPPLEMENTAL MATERIALS

The following supplemental materials are available in a separate file.

Supplemental Figure S1. Daily maximum temperature (red, °C), minimum temperature (grey, °C) and rainfall (blue, mm), from sowing to harvest date for the four yield trials. Flowering dates for 2017*_*RW not available. Flowering time of durum population is indicated by the interval between vertical black lines for each experiment.

Supplemental Figure S2. (A) Vertical change of soil moisture at different depths of soil layers, one week prior anthesis and during anthesis in the 2020_WW trial. (B) Yield of DBA Aurora and Fadda98 under different growing conditions. N represents the number of replicate under each condition. Means of the treatment were compared using Wilcoxon rank sum test.

Supplemental Figure S3. Tiller number of the 11 genotypes tracked in 2020_WW trial. GS20 and GS31 are Zadok’s scale, indicating the start of tillering and stem elongation stages, respectively. The error bars are standard deviation.

Supplemental Figure S4. Population structure of mapping population assessed by principal component analysis (PCA). (A) PCA plot of the first two components (PC1 and PC2) for 309 genotypes in the association panel. (B) Scree plot of the first ten PCs and their corresponding proportion of explained variance.

Supplemental Figure S5. Comparison of two genotypes differing in canopy closure type in the 2020 field trial. 10_82 carries four alleles associated with fast canopy closure at SNP 1095539, 3949783, 4404447 and 5324123 loci, whereas Fastoz10 carries four alleles associated with slow canopy closure at these loci. Both genotypes have similar plant density at plot level.

Supplemental Figure S6. Relationship between spike traits and grain yield in 2017_WW and 2020_WW trials. The linear regression equation, Pearson’s correlation coefficient (*R*) and significance (*p*) are displayed in each plot.

Supplemental Table S1. Durum wheat genotypes used for M × E analysis in this study.

Supplemental Table S2. Summary of results from association mapping of days to flowering (DTF), plant height (PH) and spike length (SL) in the 2020 field trial.

## DATA AVAILABILITY STATEMENT

All data supporting the findings of this study are available within the paper and within its supplementary materials published online.

