## Supplemental Figures S1-6; Supplemental Tables S1-2 for "Physiological and Genetic Drivers Underpinning Canopy Development are Associated with Durum Wheat Yield in Rainfed Environments"

Y. Kang, S.V. Haeften, S.U. Khan, E. Dinglasan, M.R. Smith, K.P. Voss-Fels, S. Alahmad and L.T. Hickey, Centre for Crop Science, Queensland Alliance for Agriculture and Food Innovation, The University of Queensland, Brisbane, QLD, Australia; D. Bustos-Korts, Biometris, Wageningen University and Research Centre, Wageningen, Netherlands; S. Vukasovic, Institute of Agronomy and Plant Breeding, Justus Liebig University Giessen, Giessen, Germany; J. Christopher and K. Chenu, Centre for Crop Science, Queensland Alliance for Agriculture and Food Innovation, The University of Queensland, Toowoomba, QLD, Australia; J.A. Able, School of Agriculture, Food & Wine, Waite Research Institute, The University of Adelaide, Urrbrae, SA, Australia; A.B. Potgieter, Centre for Crop Science, Queensland Alliance for Agriculture and Food Innovation, The University of Queensland, Gatton, QLD, Australia; D.R. Jordan and A.K. Borrell, Centre for Crop Science, Queensland Alliance for Agriculture and Food Innovation, The University of Queensland, Hermitage Research Facility, Warwick, QLD, Australia.

### Supplemental Figures

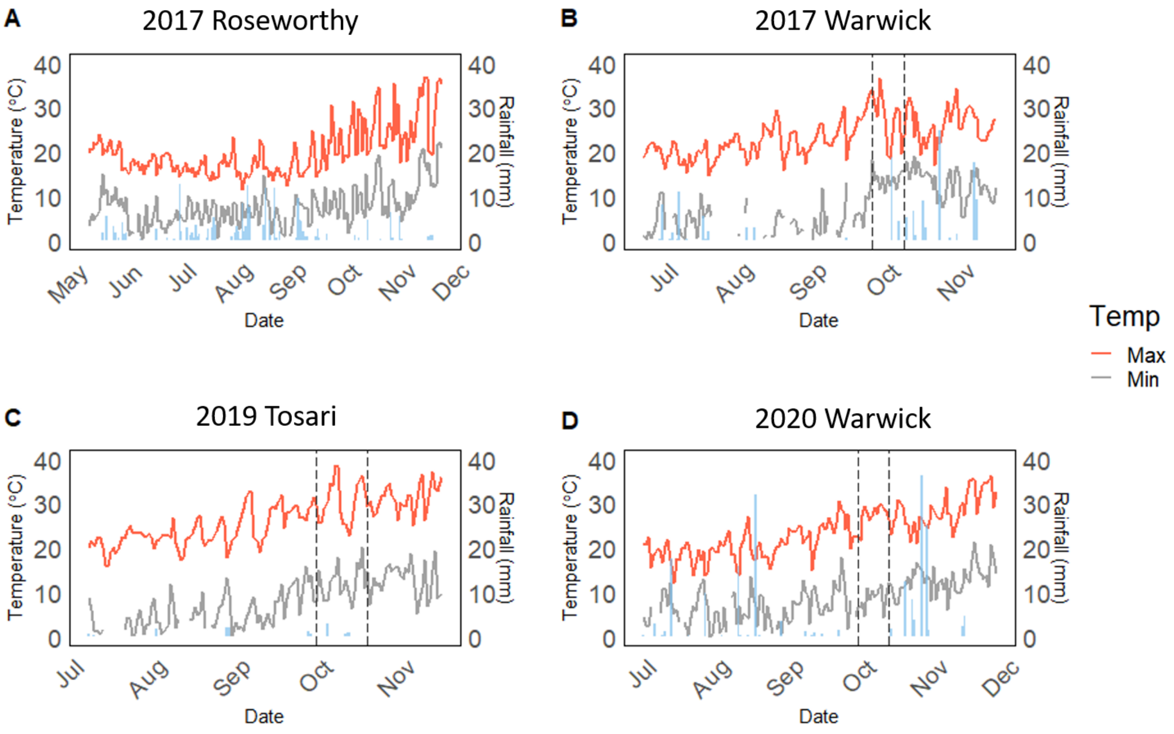


Supplemental Figure S1. Daily maximum temperature (red, °C), minimum temperature (grey, °C) and rainfall (blue, mm), from sowing to harvest date for the four yield trials. Flowering dates for 2017_RW not available. Flowering time of durum population is indicated by the interval between vertical black lines for each experiment.

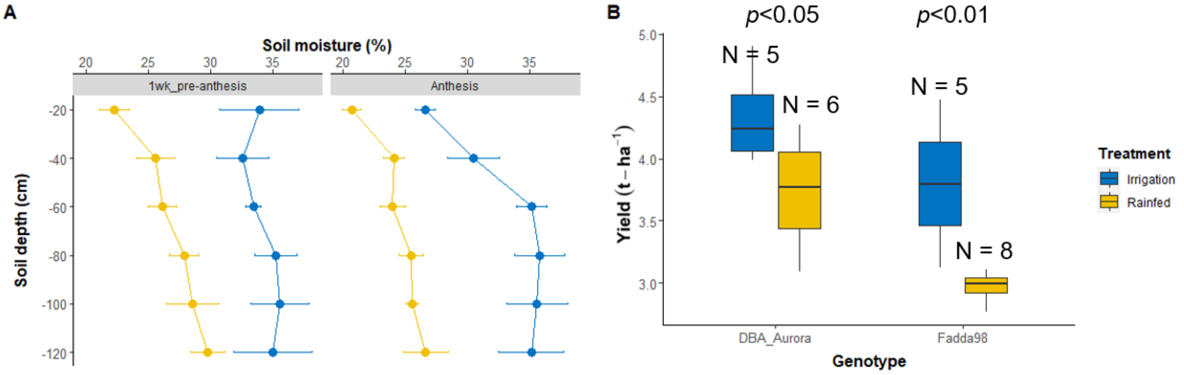


Supplemental Figure S2. (A) Vertical change of soil moisture at different depths of soil layers, one week prior anthesis and during anthesis in the 2020_WW trial. (B) Yield of DBA Aurora and Fadda98 under different growing conditions. N represents the number of replicate under each condition. Means of the treatment were compared using Wilcoxon rank sum test.

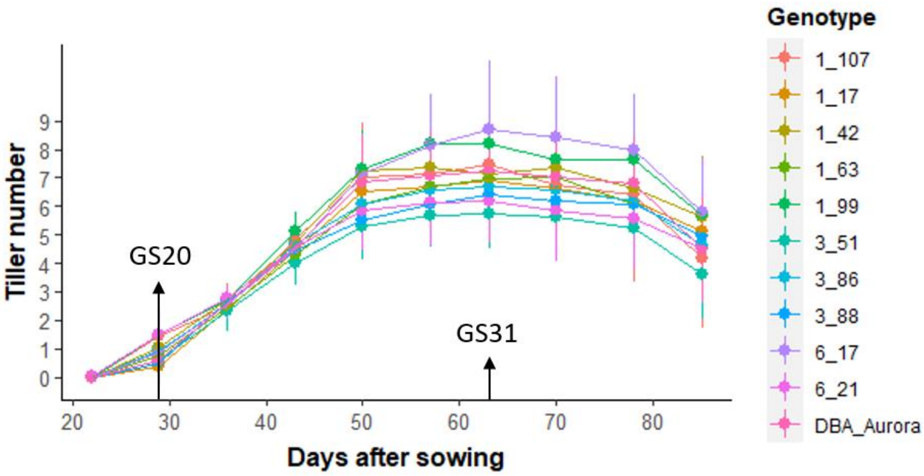


Supplemental Figure S3. Tiller number of the 11 genotypes tracked in 2020_WW trial. GS20 and GS31 are Zadok's scale, indicating the start of tillering and stem elongation stages, respectively. The error bars are standard deviation.

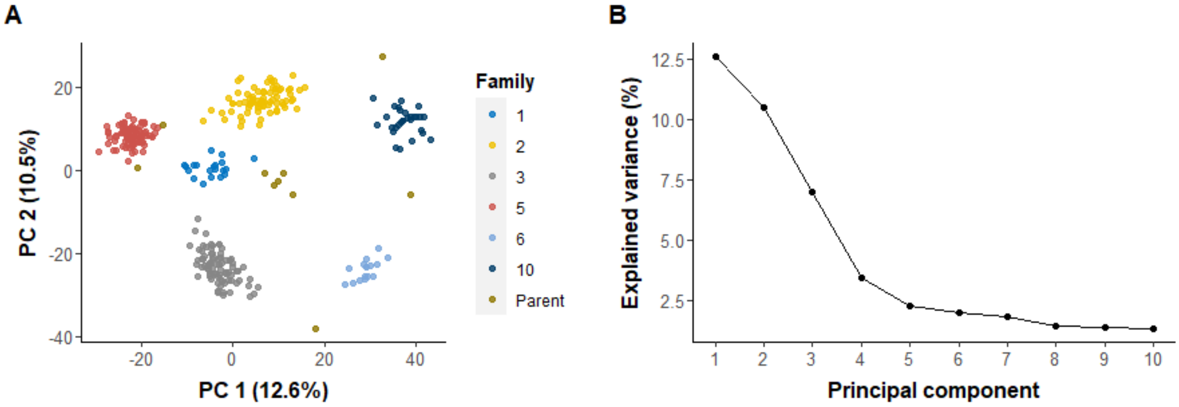


Supplemental Figure S4. Population structure of mapping population assessed by principal component analysis (PCA). (A) PCA plot of the first two components (PC1 and PC2) for 309 genotypes in the association panel. (B) Scree plot of the first ten PCs and their corresponding proportion of explained variance.

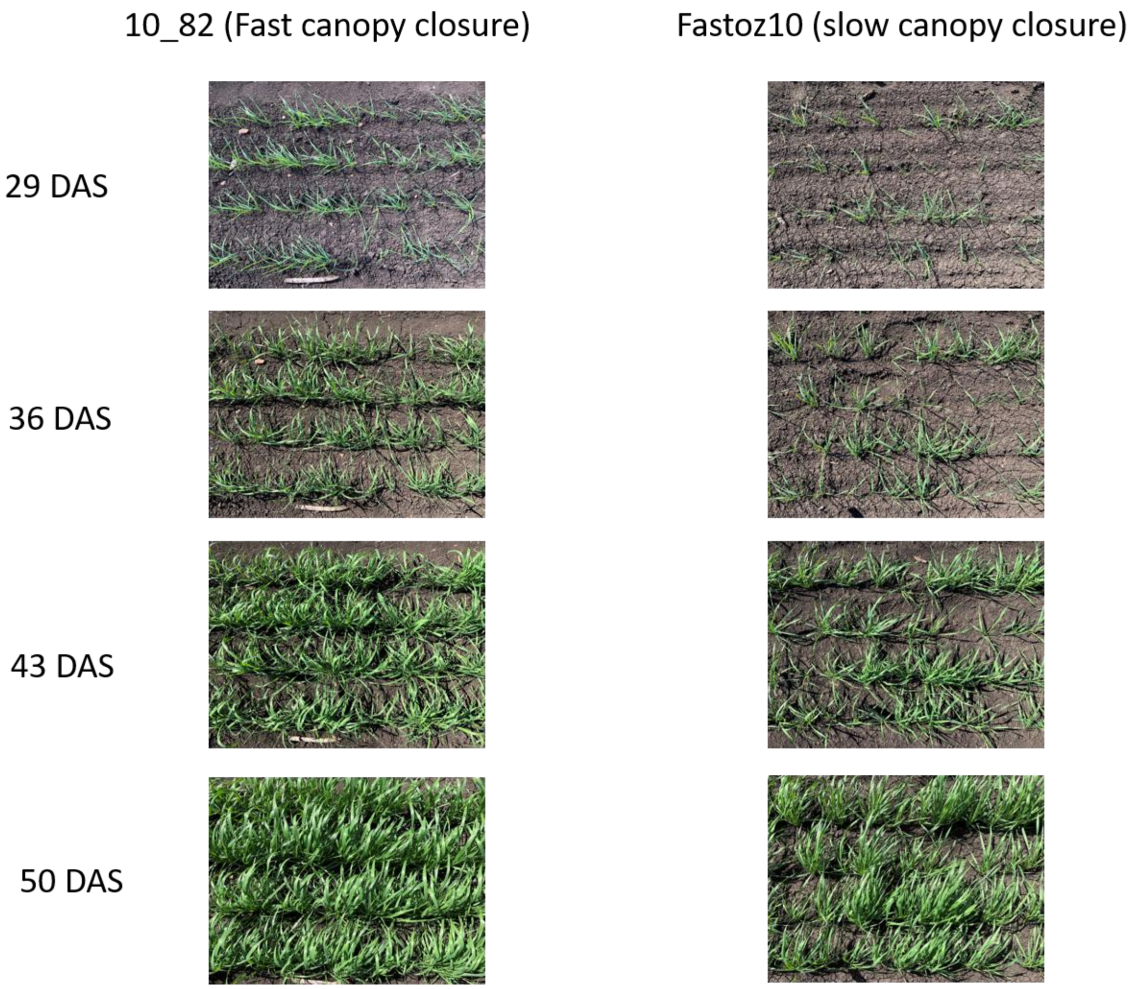


Supplemental Figure S5. Comparison of two genotypes differing in canopy closure type in the 2020 field trial. 10_82 carries four alleles associated with fast canopy closure at SNP 1095539, 3949783, 4404447 and 5324123 loci, whereas Fastoz10 carries four alleles associated with slow canopy closure at these loci. Both genotypes have similar plant density at plot level.

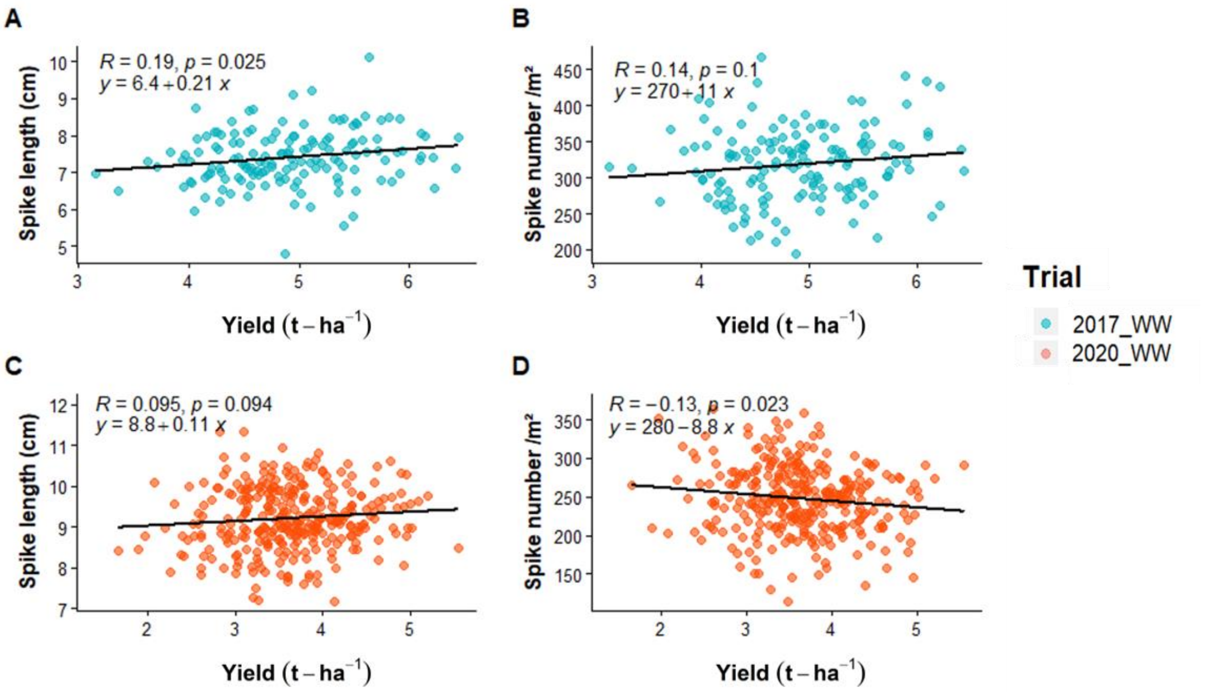


Supplemental Figure S6. Relationship between spike traits and grain yield in 2017_WW and 2020_WW trials. The linear regression equation, Pearson's correlation coefficient (*R*) and significance (*p*) are displayed in each plot.

#### Supplemental Tables

Supplemental Table S1. Durum wheat genotypes used for M × E analysis in this study.

| Genotype | Type | Pedigree | Trial |
|---|---|---|---|
| 1_103 | NAM line | DBA Aurora/Fastoz7 | 2017_RW, 2017_WW, 2019_TS, 2020_WW |
| 1_104 | NAM line | DBA Aurora/Fastoz7 | 2017_RW, 2017_WW, 2019_TS, 2020_WW |
| 1_107 | NAM line | DBA Aurora/Fastoz7 | 2017_RW, 2017_WW, 2019_TS, 2020_WW |
| 1_109 | NAM line | DBA Aurora/Fastoz7 | 2017_RW, 2017_WW, 2019_TS, 2020_WW |
| 1_128 | NAM line | DBA Aurora/Fastoz7 | 2017_RW, 2017_WW, 2019_TS, 2020_WW |
| 1_136 | NAM line | DBA Aurora/Fastoz7 | 2017_RW, 2017_WW, 2019_TS |
| 1_148 | NAM line | DBA Aurora/Fastoz7 | 2017_RW, 2017_WW, 2019_TS |
| 1_15 | NAM line | DBA Aurora/Fastoz7 | 2017_RW, 2017_WW, 2019_TS |
| 1_159 | NAM line | DBA Aurora/Fastoz7 | 2017_RW, 2017_WW, 2019_TS |
| 1_160 | NAM line | DBA Aurora/Fastoz7 | 2017_RW, 2017_WW, 2019_TS |
| 1_17 | NAM line | DBA Aurora/Fastoz7 | 2017_RW, 2017_WW, 2019_TS, 2020_WW |
| 1_2 | NAM line | DBA Aurora/Fastoz7 | 2017_RW, 2017_WW, 2019_TS, 2020_WW |
| 1_25 | NAM line | DBA Aurora/Fastoz7 | 2020_WW |
| 1_30 | NAM line | DBA Aurora/Fastoz7 | 2017_RW, 2017_WW, 2019_TS, 2020_WW |
| 1_35 | NAM line | DBA Aurora/Fastoz7 | 2017_RW, 2017_WW, 2019_TS, 2020_WW |
| 1_42 | NAM line | DBA Aurora/Fastoz7 | 2017_RW, 2017_WW, 2019_TS, 2020_WW |
| 1_48 | NAM line | DBA Aurora/Fastoz7 | 2017_RW, 2019_TS, 2020_WW |
| 1_55 | NAM line | DBA Aurora/Fastoz7 | 2017_RW, 2017_WW, 2019_TS, 2020_WW |
| 1_57 | NAM line | DBA Aurora/Fastoz7 | 2017_RW, 2017_WW, 2019_TS, 2020_WW |
| 1_63 | NAM line | DBA Aurora/Fastoz7 | 2017_RW, 2017_WW, 2019_TS, 2020_WW |
| 1_70 | NAM line | DBA Aurora/Fastoz7 | 2017_RW, 2017_WW, 2019_TS, 2020_WW |
| 1_77 | NAM line | DBA Aurora/Fastoz7 | 2017_RW, 2017_WW, 2019_TS, 2020_WW |
| 1_83 | NAM line | DBA Aurora/Fastoz7 | 2017_RW, 2017_WW, 2019_TS, 2020_WW |

| | | | |
|---|---|---|---|
| 1_99 | NAM line | DBA Aurora/Fastoz7 | 2017_RW, 2017_WW, 2019_TS, 2020_WW |
| 10_1 | NAM line | Jandaroi/Outrob4 | 2020_WW |
| 10_100 | NAM line | Jandaroi/Outrob4 | 2017_RW, 2017_WW, 2019_TS |
| 10_104 | NAM line | Jandaroi/Outrob4 | 2020_WW |
| 10_106 | NAM line | Jandaroi/Outrob4 | 2020_WW |
| 10_124 | NAM line | Jandaroi/Outrob4 | 2017_RW, 2017_WW, 2019_TS, 2020_WW |
| 10_130 | NAM line | Jandaroi/Outrob4 | 2017_RW, 2017_WW, 2019_TS |
| 10_14 | NAM line | Jandaroi/Outrob4 | 2017_RW, 2017_WW, 2019_TS |
| 10_144 | NAM line | Jandaroi/Outrob4 | 2017_RW, 2017_WW, 2019_TS |
| 10_148 | NAM line | Jandaroi/Outrob4 | 2017_RW, 2017_WW |
| 10_17 | NAM line | Jandaroi/Outrob4 | 2020_WW |
| 10_2 | NAM line | Jandaroi/Outrob4 | 2020_WW |
| 10_20 | NAM line | Jandaroi/Outrob4 | 2017_RW, 2017_WW, 2019_TS, 2020_WW |
| 10_25 | NAM line | Jandaroi/Outrob4 | 2020_WW |
| 10_29 | NAM line | Jandaroi/Outrob4 | 2017_RW, 2017_WW, 2019_TS, 2020_WW |
| 10_3 | NAM line | Jandaroi/Outrob4 | 2020_WW |
| 10_31 | NAM line | Jandaroi/Outrob4 | 2020_WW |
| 10_35 | NAM line | Jandaroi/Outrob4 | 2020_WW |
| 10_36 | NAM line | Jandaroi/Outrob4 | 2020_WW |
| 10_38 | NAM line | Jandaroi/Outrob4 | 2020_WW |
| 10_44 | NAM line | Jandaroi/Outrob4 | 2020_WW |
| 10_53 | NAM line | Jandaroi/Outrob4 | 2020_WW |
| 10_54 | NAM line | Jandaroi/Outrob4 | 2020_WW |
| 10_56 | NAM line | Jandaroi/Outrob4 | 2017_RW, 2017_WW, 2019_TS |
| 10_6 | NAM line | Jandaroi/Outrob4 | 2017_RW, 2017_WW, 2019_TS, 2020_WW |
| 10_61 | NAM line | Jandaroi/Outrob4 | 2017_RW, 2017_WW, 2019_TS |
| 10_63 | NAM line | Jandaroi/Outrob4 | 2020_WW |
| 10_67 | NAM line | Jandaroi/Outrob4 | 2017_RW, 2017_WW, 2019_TS, 2020_WW |
| 10_68 | NAM line | Jandaroi/Outrob4 | 2020_WW |

| | | | |
|---|---|---|---|
| 10_69 | NAM line | Jandaroi/Outrob4 | 2020_WW |
| 10_7 | NAM line | Jandaroi/Outrob4 | 2020_WW |
| 10_70 | NAM line | Jandaroi/Outrob4 | 2020_WW |
| 10_72 | NAM line | Jandaroi/Outrob4 | 2017_RW, 2017_WW, 2019_TS, 2020_WW |
| 10_74 | NAM line | Jandaroi/Outrob4 | 2020_WW |
| 10_76 | NAM line | Jandaroi/Outrob4 | 2017_RW, 2017_WW, 2019_TS |
| 10_77 | NAM line | Jandaroi/Outrob4 | 2020_WW |
| 10_81 | NAM line | Jandaroi/Outrob4 | 2017_RW, 2017_WW, 2019_TS |
| 10_82 | NAM line | Jandaroi/Outrob4 | 2020_WW |
| 10_88 | NAM line | Jandaroi/Outrob4 | 2017_RW, 2017_WW, 2019_TS |
| 10_90 | NAM line | Jandaroi/Outrob4 | 2020_WW |
| 10_95 | NAM line | Jandaroi/Outrob4 | 2017_RW, 2017_WW, 2019_TS |
| 2_1 | NAM line | DBA Aurora/Outrob4 | 2020_WW |
| 2_10 | NAM line | DBA Aurora/Outrob4 | 2020_WW |
| 2_103 | NAM line | DBA Aurora/Outrob4 | 2017_RW, 2017_WW, 2019_TS, 2020_WW |
| 2_104 | NAM line | DBA Aurora/Outrob4 | 2020_WW |
| 2_107 | NAM line | DBA Aurora/Outrob4 | 2017_RW, 2017_WW, 2019_TS, 2020_WW |
| 2_109 | NAM line | DBA Aurora/Outrob4 | 2020_WW |
| 2_110 | NAM line | DBA Aurora/Outrob4 | 2020_WW |
| 2_114 | NAM line | DBA Aurora/Outrob4 | 2017_RW, 2017_WW, 2019_TS, 2020_WW |
| 2_12 | NAM line | DBA Aurora/Outrob4 | 2020_WW |
| 2_13 | NAM line | DBA Aurora/Outrob4 | 2020_WW |
| 2_14 | NAM line | DBA Aurora/Outrob4 | 2020_WW |
| 2_15 | NAM line | DBA Aurora/Outrob4 | 2020_WW |
| 2_16 | NAM line | DBA Aurora/Outrob4 | 2020_WW |
| 2_17 | NAM line | DBA Aurora/Outrob4 | 2020_WW |
| 2_19 | NAM line | DBA Aurora/Outrob4 | 2020_WW |
| 2_2 | NAM line | DBA Aurora/Outrob4 | 2020_WW |
| 2_20 | NAM line | DBA Aurora/Outrob4 | 2020_WW |

| | | | |
|---|---|---|---|
| 2_21 | NAM line | DBA Aurora/Outrob4 | 2020_WW |
| 2_22 | NAM line | DBA Aurora/Outrob4 | 2020_WW |
| 2_23 | NAM line | DBA Aurora/Outrob4 | 2020_WW |
| 2_24 | NAM line | DBA Aurora/Outrob4 | 2020_WW |
| 2_26 | NAM line | DBA Aurora/Outrob4 | 2020_WW |
| 2_29 | NAM line | DBA Aurora/Outrob4 | 2020_WW |
| 2_30 | NAM line | DBA Aurora/Outrob4 | 2020_WW |
| 2_31 | NAM line | DBA Aurora/Outrob4 | 2020_WW |
| 2_35 | NAM line | DBA Aurora/Outrob4 | 2020_WW |
| 2_36 | NAM line | DBA Aurora/Outrob4 | 2020_WW |
| 2_37 | NAM line | DBA Aurora/Outrob4 | 2020_WW |
| 2_38 | NAM line | DBA Aurora/Outrob4 | 2020_WW |
| 2_39 | NAM line | DBA Aurora/Outrob4 | 2020_WW |
| 2_4 | NAM line | DBA Aurora/Outrob4 | 2017_RW, 2017_WW, 2019_TS, 2020_WW |
| 2_41 | NAM line | DBA Aurora/Outrob4 | 2017_RW, 2017_WW, 2019_TS, 2020_WW |
| 2_43 | NAM line | DBA Aurora/Outrob4 | 2020_WW |
| 2_44 | NAM line | DBA Aurora/Outrob4 | 2020_WW |
| 2_45 | NAM line | DBA Aurora/Outrob4 | 2020_WW |
| 2_46 | NAM line | DBA Aurora/Outrob4 | 2020_WW |
| 2_48 | NAM line | DBA Aurora/Outrob4 | 2020_WW |
| 2_49 | NAM line | DBA Aurora/Outrob4 | 2020_WW |
| 2_5 | NAM line | DBA Aurora/Outrob4 | 2020_WW |
| 2_50 | NAM line | DBA Aurora/Outrob4 | 2020_WW |
| 2_51 | NAM line | DBA Aurora/Outrob4 | 2020_WW |
| 2_54 | NAM line | DBA Aurora/Outrob4 | 2020_WW |
| 2_56 | NAM line | DBA Aurora/Outrob4 | 2017_RW, 2017_WW, 2019_TS, 2020_WW |
| 2_59 | NAM line | DBA Aurora/Outrob4 | 2020_WW |
| 2_6 | NAM line | DBA Aurora/Outrob4 | 2020_WW |
| 2_61 | NAM line | DBA Aurora/Outrob4 | 2020_WW |

| | | | |
|---|---|---|---|
| 2_62 | NAM line | DBA Aurora/Outrob4 | 2020_WW |
| 2_64 | NAM line | DBA Aurora/Outrob4 | 2020_WW |
| 2_66 | NAM line | DBA Aurora/Outrob4 | 2020_WW |
| 2_67 | NAM line | DBA Aurora/Outrob4 | 2020_WW |
| 2_68 | NAM line | DBA Aurora/Outrob4 | 2020_WW |
| 2_69 | NAM line | DBA Aurora/Outrob4 | 2020_WW |
| 2_7 | NAM line | DBA Aurora/Outrob4 | 2020_WW |
| 2_70 | NAM line | DBA Aurora/Outrob4 | 2020_WW |
| 2_71 | NAM line | DBA Aurora/Outrob4 | 2020_WW |
| 2_72 | NAM line | DBA Aurora/Outrob4 | 2020_WW |
| 2_73 | NAM line | DBA Aurora/Outrob4 | 2020_WW |
| 2_74 | NAM line | DBA Aurora/Outrob4 | 2020_WW |
| 2_75 | NAM line | DBA Aurora/Outrob4 | 2020_WW |
| 2_76 | NAM line | DBA Aurora/Outrob4 | 2020_WW |
| 2_77 | NAM line | DBA Aurora/Outrob4 | 2020_WW |
| 2_78 | NAM line | DBA Aurora/Outrob4 | 2017_RW, 2019_TS |
| 2_8 | NAM line | DBA Aurora/Outrob4 | 2020_WW |
| 2_82 | NAM line | DBA Aurora/Outrob4 | 2020_WW |
| 2_84 | NAM line | DBA Aurora/Outrob4 | 2020_WW |
| 2_85 | NAM line | DBA Aurora/Outrob4 | 2020_WW |
| 2_86 | NAM line | DBA Aurora/Outrob4 | 2017_RW, 2017_WW, 2019_TS, 2020_WW |
| 2_87 | NAM line | DBA Aurora/Outrob4 | 2020_WW |
| 2_88 | NAM line | DBA Aurora/Outrob4 | 2020_WW |
| 2_9 | NAM line | DBA Aurora/Outrob4 | 2020_WW |
| 2_91 | NAM line | DBA Aurora/Outrob4 | 2017_RW, 2017_WW, 2019_TS, 2020_WW |
| 2_93 | NAM line | DBA Aurora/Outrob4 | 2020_WW |
| 2_94 | NAM line | DBA Aurora/Outrob4 | 2020_WW |
| 2_95 | NAM line | DBA Aurora/Outrob4 | 2020_WW |
| 2_98 | NAM line | DBA Aurora/Outrob4 | 2017_RW, 2017_WW, 2019_TS, 2020_WW |

| | | | |
|---|---|---|---|
| 3_10 | NAM line | DBA Aurora/Fastoz8 | 2019_TS, 2020_WW |
| 3_102 | NAM line | DBA Aurora/Fastoz8 | 2019_TS, 2020_WW |
| 3_103 | NAM line | DBA Aurora/Fastoz8 | 2019_TS, 2020_WW |
| 3_105 | NAM line | DBA Aurora/Fastoz8 | 2019_TS, 2020_WW |
| 3_12 | NAM line | DBA Aurora/Fastoz8 | 2017_RW, 2017_WW, 2019_TS, 2020_WW |
| 3_13 | NAM line | DBA Aurora/Fastoz8 | 2019_TS, 2020_WW |
| 3_130 | NAM line | DBA Aurora/Fastoz8 | 2017_RW, 2017_WW, 2019_TS |
| 3_132 | NAM line | DBA Aurora/Fastoz8 | 2017_RW, 2017_WW, 2019_TS, 2020_WW |
| 3_15 | NAM line | DBA Aurora/Fastoz8 | 2019_TS |
| 3_17 | NAM line | DBA Aurora/Fastoz8 | 2019_TS, 2020_WW |
| 3_18 | NAM line | DBA Aurora/Fastoz8 | 2019_TS, 2020_WW |
| 3_19 | NAM line | DBA Aurora/Fastoz8 | 2017_RW, 2017_WW, 2019_TS, 2020_WW |
| 3_2 | NAM line | DBA Aurora/Fastoz8 | 2020_WW |
| 3_21 | NAM line | DBA Aurora/Fastoz8 | 2019_TS, 2020_WW |
| 3_22 | NAM line | DBA Aurora/Fastoz8 | 2019_TS, 2020_WW |
| 3_23 | NAM line | DBA Aurora/Fastoz8 | 2019_TS, 2020_WW |
| 3_24 | NAM line | DBA Aurora/Fastoz8 | 2019_TS, 2020_WW |
| 3_25 | NAM line | DBA Aurora/Fastoz8 | 2019_TS, 2020_WW |
| 3_26 | NAM line | DBA Aurora/Fastoz8 | 2019_TS, 2020_WW |
| 3_27 | NAM line | DBA Aurora/Fastoz8 | 2019_TS, 2020_WW |
| 3_28 | NAM line | DBA Aurora/Fastoz8 | 2019_TS, 2020_WW |
| 3_29 | NAM line | DBA Aurora/Fastoz8 | 2019_TS, 2020_WW |
| 3_30 | NAM line | DBA Aurora/Fastoz8 | 2017_RW, 2017_WW, 2019_TS, 2020_WW |
| 3_31 | NAM line | DBA Aurora/Fastoz8 | 2019_TS |
| 3_32 | NAM line | DBA Aurora/Fastoz8 | 2019_TS, 2020_WW |
| 3_33 | NAM line | DBA Aurora/Fastoz8 | 2019_TS, 2020_WW |
| 3_34 | NAM line | DBA Aurora/Fastoz8 | 2019_TS, 2020_WW |
| 3_35 | NAM line | DBA Aurora/Fastoz8 | 2019_TS, 2020_WW |
| 3_36 | NAM line | DBA Aurora/Fastoz8 | 2019_TS, 2020_WW |

| | | | |
|---|---|---|---|
| 3_37 | NAM line | DBA Aurora/Fastoz8 | 2019_TS, 2020_WW |
| 3_38 | NAM line | DBA Aurora/Fastoz8 | 2019_TS, 2020_WW |
| 3_39 | NAM line | DBA Aurora/Fastoz8 | 2019_TS, 2020_WW |
| 3_4 | NAM line | DBA Aurora/Fastoz8 | 2019_TS, 2020_WW |
| 3_40 | NAM line | DBA Aurora/Fastoz8 | 2017_RW, 2017_WW, 2019_TS, 2020_WW |
| 3_41 | NAM line | DBA Aurora/Fastoz8 | 2019_TS, 2020_WW |
| 3_42 | NAM line | DBA Aurora/Fastoz8 | 2019_TS, 2020_WW |
| 3_43 | NAM line | DBA Aurora/Fastoz8 | 2019_TS, 2020_WW |
| 3_44 | NAM line | DBA Aurora/Fastoz8 | 2019_TS, 2020_WW |
| 3_45 | NAM line | DBA Aurora/Fastoz8 | 2019_TS, 2020_WW |
| 3_46 | NAM line | DBA Aurora/Fastoz8 | 2019_TS, 2020_WW |
| 3_48 | NAM line | DBA Aurora/Fastoz8 | 2019_TS, 2020_WW |
| 3_49 | NAM line | DBA Aurora/Fastoz8 | 2019_TS |
| 3_5 | NAM line | DBA Aurora/Fastoz8 | 2017_RW, 2017_WW, 2019_TS, 2020_WW |
| 3_50 | NAM line | DBA Aurora/Fastoz8 | 2017_RW, 2017_WW, 2019_TS, 2020_WW |
| 3_51 | NAM line | DBA Aurora/Fastoz8 | 2019_TS, 2020_WW |
| 3_52 | NAM line | DBA Aurora/Fastoz8 | 2019_TS, 2020_WW |
| 3_54 | NAM line | DBA Aurora/Fastoz8 | 2019_TS, 2020_WW |
| 3_55 | NAM line | DBA Aurora/Fastoz8 | 2019_TS, 2020_WW |
| 3_56 | NAM line | DBA Aurora/Fastoz8 | 2019_TS, 2020_WW |
| 3_57 | NAM line | DBA Aurora/Fastoz8 | 2019_TS, 2020_WW |
| 3_58 | NAM line | DBA Aurora/Fastoz8 | 2019_TS |
| 3_59 | NAM line | DBA Aurora/Fastoz8 | 2019_TS, 2020_WW |
| 3_6 | NAM line | DBA Aurora/Fastoz8 | 2019_TS |
| 3_60 | NAM line | DBA Aurora/Fastoz8 | 2017_RW, 2017_WW, 2019_TS, 2020_WW |
| 3_61 | NAM line | DBA Aurora/Fastoz8 | 2019_TS, 2020_WW |
| 3_62 | NAM line | DBA Aurora/Fastoz8 | 2019_TS, 2020_WW |
| 3_63 | NAM line | DBA Aurora/Fastoz8 | 2019_TS, 2020_WW |
| 3_64 | NAM line | DBA Aurora/Fastoz8 | 2019_TS, 2020_WW |

| | | | |
|---|---|---|---|
| 3_65 | NAM line | DBA Aurora/Fastoz8 | 2019_TS |
| 3_68 | NAM line | DBA Aurora/Fastoz8 | 2019_TS, 2020_WW |
| 3_69 | NAM line | DBA Aurora/Fastoz8 | 2019_TS, 2020_WW |
| 3_70 | NAM line | DBA Aurora/Fastoz8 | 2019_TS, 2020_WW |
| 3_71 | NAM line | DBA Aurora/Fastoz8 | 2019_TS, 2020_WW |
| 3_72 | NAM line | DBA Aurora/Fastoz8 | 2019_TS, 2020_WW |
| 3_74 | NAM line | DBA Aurora/Fastoz8 | 2017_RW, 2017_WW, 2019_TS, 2020_WW |
| 3_75 | NAM line | DBA Aurora/Fastoz8 | 2019_TS, 2020_WW |
| 3_76 | NAM line | DBA Aurora/Fastoz8 | 2019_TS, 2020_WW |
| 3_78 | NAM line | DBA Aurora/Fastoz8 | 2019_TS |
| 3_79 | NAM line | DBA Aurora/Fastoz8 | 2019_TS, 2020_WW |
| 3_80 | NAM line | DBA Aurora/Fastoz8 | 2019_TS, 2020_WW |
| 3_81 | NAM line | DBA Aurora/Fastoz8 | 2019_TS, 2020_WW |
| 3_82 | NAM line | DBA Aurora/Fastoz8 | 2019_TS, 2020_WW |
| 3_83 | NAM line | DBA Aurora/Fastoz8 | 2019_TS |
| 3_84 | NAM line | DBA Aurora/Fastoz8 | 2019_TS, 2020_WW |
| 3_85 | NAM line | DBA Aurora/Fastoz8 | 2019_TS, 2020_WW |
| 3_86 | NAM line | DBA Aurora/Fastoz8 | 2019_TS, 2020_WW |
| 3_87 | NAM line | DBA Aurora/Fastoz8 | 2019_TS, 2020_WW |
| 3_88 | NAM line | DBA Aurora/Fastoz8 | 2019_TS, 2020_WW |
| 3_89 | NAM line | DBA Aurora/Fastoz8 | 2019_TS, 2020_WW |
| 3_9 | NAM line | DBA Aurora/Fastoz8 | 2019_TS, 2020_WW |
| 3_90 | NAM line | DBA Aurora/Fastoz8 | 2019_TS, 2020_WW |
| 3_91 | NAM line | DBA Aurora/Fastoz8 | 2019_TS |
| 3_92 | NAM line | DBA Aurora/Fastoz8 | 2019_TS, 2020_WW |
| 3_93 | NAM line | DBA Aurora/Fastoz8 | 2019_TS, 2020_WW |
| 3_94 | NAM line | DBA Aurora/Fastoz8 | 2019_TS, 2020_WW |
| 3_95 | NAM line | DBA Aurora/Fastoz8 | 2019_TS, 2020_WW |
| 3_96 | NAM line | DBA Aurora/Fastoz8 | 2019_TS, 2020_WW |

| | | | |
|---|---|---|---|
| 3_97 | NAM line | DBA Aurora/Fastoz8 | 2019_TS, 2020_WW |
| 3_98 | NAM line | DBA Aurora/Fastoz8 | 2019_TS, 2020_WW |
| 3_99 | NAM line | DBA Aurora/Fastoz8 | 2019_TS, 2020_WW |
| 4_111 | NAM line | DBA Aurora/Fadda98 | 2017_RW, 2017_WW, 2019_TS |
| 4_123 | NAM line | DBA Aurora/Fadda98 | 2017_RW, 2017_WW, 2019_TS |
| 4_4 | NAM line | DBA Aurora/Fadda98 | 2017_RW, 2017_WW, 2019_TS |
| 4_49 | NAM line | DBA Aurora/Fadda98 | 2017_RW, 2017_WW, 2019_TS |
| 4_61 | NAM line | DBA Aurora/Fadda98 | 2017_RW, 2017_WW, 2019_TS |
| 5_1 | NAM line | DBA Aurora/Fastoz3 | 2020_WW |
| 5_10 | NAM line | DBA Aurora/Fastoz3 | 2020_WW |
| 5_101 | NAM line | DBA Aurora/Fastoz3 | 2020_WW |
| 5_102 | NAM line | DBA Aurora/Fastoz3 | 2020_WW |
| 5_104 | NAM line | DBA Aurora/Fastoz3 | 2020_WW |
| 5_105 | NAM line | DBA Aurora/Fastoz3 | 2020_WW |
| 5_106 | NAM line | DBA Aurora/Fastoz3 | 2020_WW |
| 5_107 | NAM line | DBA Aurora/Fastoz3 | 2017_RW, 2017_WW, 2019_TS |
| 5_109 | NAM line | DBA Aurora/Fastoz3 | 2020_WW |
| 5_11 | NAM line | DBA Aurora/Fastoz3 | 2020_WW |
| 5_110 | NAM line | DBA Aurora/Fastoz3 | 2020_WW |
| 5_111 | NAM line | DBA Aurora/Fastoz3 | 2020_WW |
| 5_112 | NAM line | DBA Aurora/Fastoz3 | 2017_RW, 2017_WW, 2019_TS, 2020_WW |
| 5_113 | NAM line | DBA Aurora/Fastoz3 | 2020_WW |
| 5_114 | NAM line | DBA Aurora/Fastoz3 | 2020_WW |
| 5_115 | NAM line | DBA Aurora/Fastoz3 | 2020_WW |
| 5_13 | NAM line | DBA Aurora/Fastoz3 | 2020_WW |
| 5_138 | NAM line | DBA Aurora/Fastoz3 | 2017_RW, 2017_WW, 2019_TS, 2020_WW |
| 5_140 | NAM line | DBA Aurora/Fastoz3 | 2017_RW, 2017_WW, 2019_TS, 2020_WW |
| 5_15 | NAM line | DBA Aurora/Fastoz3 | 2020_WW |
| 5_152 | NAM line | DBA Aurora/Fastoz3 | 2017_RW, 2017_WW, 2019_TS, 2020_WW |

| | | | |
|---|---|---|---|
| 5_17 | NAM line | DBA Aurora/Fastoz3 | 2020_WW |
| 5_18 | NAM line | DBA Aurora/Fastoz3 | 2020_WW |
| 5_19 | NAM line | DBA Aurora/Fastoz3 | 2020_WW |
| 5_20 | NAM line | DBA Aurora/Fastoz3 | 2020_WW |
| 5_21 | NAM line | DBA Aurora/Fastoz3 | 2020_WW |
| 5_22 | NAM line | DBA Aurora/Fastoz3 | 2020_WW |
| 5_23 | NAM line | DBA Aurora/Fastoz3 | 2020_WW |
| 5_27 | NAM line | DBA Aurora/Fastoz3 | 2020_WW |
| 5_28 | NAM line | DBA Aurora/Fastoz3 | 2020_WW |
| 5_29 | NAM line | DBA Aurora/Fastoz3 | 2020_WW |
| 5_3 | NAM line | DBA Aurora/Fastoz3 | 2020_WW |
| 5_30 | NAM line | DBA Aurora/Fastoz3 | 2020_WW |
| 5_31 | NAM line | DBA Aurora/Fastoz3 | 2020_WW |
| 5_33 | NAM line | DBA Aurora/Fastoz3 | 2020_WW |
| 5_35 | NAM line | DBA Aurora/Fastoz3 | 2020_WW |
| 5_36 | NAM line | DBA Aurora/Fastoz3 | 2020_WW |
| 5_37 | NAM line | DBA Aurora/Fastoz3 | 2020_WW |
| 5_38 | NAM line | DBA Aurora/Fastoz3 | 2020_WW |
| 5_39 | NAM line | DBA Aurora/Fastoz3 | 2020_WW |
| 5_40 | NAM line | DBA Aurora/Fastoz3 | 2017_RW, 2019_TS |
| 5_42 | NAM line | DBA Aurora/Fastoz3 | 2020_WW |
| 5_43 | NAM line | DBA Aurora/Fastoz3 | 2020_WW |
| 5_44 | NAM line | DBA Aurora/Fastoz3 | 2020_WW |
| 5_45 | NAM line | DBA Aurora/Fastoz3 | 2020_WW |
| 5_46 | NAM line | DBA Aurora/Fastoz3 | 2020_WW |
| 5_47 | NAM line | DBA Aurora/Fastoz3 | 2020_WW |
| 5_48 | NAM line | DBA Aurora/Fastoz3 | 2020_WW |
| 5_49 | NAM line | DBA Aurora/Fastoz3 | 2020_WW |
| 5_5 | NAM line | DBA Aurora/Fastoz3 | 2020_WW |

| | | | |
|---|---|---|---|
| 5_50 | NAM line | DBA Aurora/Fastoz3 | 2020_WW |
| 5_51 | NAM line | DBA Aurora/Fastoz3 | 2017_RW, 2017_WW, 2019_TS, 2020_WW |
| 5_53 | NAM line | DBA Aurora/Fastoz3 | 2020_WW |
| 5_55 | NAM line | DBA Aurora/Fastoz3 | 2020_WW |
| 5_56 | NAM line | DBA Aurora/Fastoz3 | 2020_WW |
| 5_57 | NAM line | DBA Aurora/Fastoz3 | 2020_WW |
| 5_59 | NAM line | DBA Aurora/Fastoz3 | 2020_WW |
| 5_6 | NAM line | DBA Aurora/Fastoz3 | 2020_WW |
| 5_60 | NAM line | DBA Aurora/Fastoz3 | 2017_RW, 2017_WW, 2019_TS, 2020_WW |
| 5_61 | NAM line | DBA Aurora/Fastoz3 | 2020_WW |
| 5_64 | NAM line | DBA Aurora/Fastoz3 | 2017_RW, 2017_WW, 2019_TS, 2020_WW |
| 5_65 | NAM line | DBA Aurora/Fastoz3 | 2020_WW |
| 5_67 | NAM line | DBA Aurora/Fastoz3 | 2020_WW |
| 5_69 | NAM line | DBA Aurora/Fastoz3 | 2020_WW |
| 5_7 | NAM line | DBA Aurora/Fastoz3 | 2020_WW |
| 5_70 | NAM line | DBA Aurora/Fastoz3 | 2020_WW |
| 5_71 | NAM line | DBA Aurora/Fastoz3 | 2020_WW |
| 5_73 | NAM line | DBA Aurora/Fastoz3 | 2020_WW |
| 5_75 | NAM line | DBA Aurora/Fastoz3 | 2020_WW |
| 5_76 | NAM line | DBA Aurora/Fastoz3 | 2020_WW |
| 5_78 | NAM line | DBA Aurora/Fastoz3 | 2020_WW |
| 5_79 | NAM line | DBA Aurora/Fastoz3 | 2020_WW |
| 5_8 | NAM line | DBA Aurora/Fastoz3 | 2020_WW |
| 5_80 | NAM line | DBA Aurora/Fastoz3 | 2020_WW |
| 5_85 | NAM line | DBA Aurora/Fastoz3 | 2020_WW |
| 5_86 | NAM line | DBA Aurora/Fastoz3 | 2020_WW |
| 5_88 | NAM line | DBA Aurora/Fastoz3 | 2020_WW |
| 5_9 | NAM line | DBA Aurora/Fastoz3 | 2020_WW |
| 5_90 | NAM line | DBA Aurora/Fastoz3 | 2020_WW |

| | | | |
|---|---|---|---|
| 5_94 | NAM line | DBA Aurora/Fastoz3 | 2020_WW |
| 5_97 | NAM line | DBA Aurora/Fastoz3 | 2020_WW |
| 5_99 | NAM line | DBA Aurora/Fastoz3 | 2020_WW |
| 6_101 | NAM line | Jandaroi/Fastoz8 | 2017_RW, 2017_WW |
| 6_116 | NAM line | Jandaroi/Fastoz8 | 2017_RW, 2017_WW, 2020_WW |
| 6_142 | NAM line | Jandaroi/Fastoz8 | 2017_RW, 2017_WW |
| 6_143 | NAM line | Jandaroi/Fastoz8 | 2017_RW, 2017_WW, 2020_WW |
| 6_148 | NAM line | Jandaroi/Fastoz8 | 2017_RW, 2017_WW, 2020_WW |
| 6_151 | NAM line | Jandaroi/Fastoz8 | 2017_RW, 2017_WW, 2020_WW |
| 6_156 | NAM line | Jandaroi/Fastoz8 | 2017_RW, 2017_WW, 2019_TS, 2020_WW |
| 6_17 | NAM line | Jandaroi/Fastoz8 | 2017_RW, 2017_WW, 2019_TS, 2020_WW |
| 6_21 | NAM line | Jandaroi/Fastoz8 | 2017_RW, 2017_WW, 2019_TS, 2020_WW |
| 6_34 | NAM line | Jandaroi/Fastoz8 | 2017_RW, 2017_WW, 2019_TS, 2020_WW |
| 6_38 | NAM line | Jandaroi/Fastoz8 | 2017_RW, 2017_WW, 2019_TS, 2020_WW |
| 6_49 | NAM line | Jandaroi/Fastoz8 | 2017_RW, 2017_WW, 2019_TS, 2020_WW |
| 6_53 | NAM line | Jandaroi/Fastoz8 | 2017_RW, 2017_WW, 2019_TS, 2020_WW |
| 6_62 | NAM line | Jandaroi/Fastoz8 | 2017_RW, 2017_WW, 2019_TS, 2020_WW |
| 6_64 | NAM line | Jandaroi/Fastoz8 | 2017_RW, 2017_WW |
| 6_70 | NAM line | Jandaroi/Fastoz8 | 2017_RW, 2017_WW, 2019_TS, 2020_WW |
| 6_79 | NAM line | Jandaroi/Fastoz8 | 2017_RW, 2017_WW, 2019_TS |
| 6_86 | NAM line | Jandaroi/Fastoz8 | 2017_RW, 2017_WW, 2019_TS, 2020_WW |
| 6_9 | NAM line | Jandaroi/Fastoz8 | 2017_RW, 2017_WW, 2019_TS, 2020_WW |
| 6_98 | NAM line | Jandaroi/Fastoz8 | 2017_RW, 2017_WW, 2019_TS, 2020_WW |
| 7_101 | NAM line | Jandaroi/Fastoz10 | 2017_RW, 2017_WW, 2019_TS |
| 7_145 | NAM line | Jandaroi/Fastoz10 | 2017_RW, 2017_WW, 2019_TS |
| 7_15 | NAM line | Jandaroi/Fastoz10 | 2017_RW, 2017_WW, 2019_TS |
| 7_19 | NAM line | Jandaroi/Fastoz10 | 2017_RW, 2017_WW, 2019_TS |
| 7_34 | NAM line | Jandaroi/Fastoz10 | 2017_RW, 2017_WW, 2019_TS |
| 7_42 | NAM line | Jandaroi/Fastoz10 | 2017_RW, 2017_WW, 2019_TS |

| | | | |
|---|---|---|---|
| 7_57 | NAM line | Jandaroi/Fastoz10 | 2017_RW, 2017_WW, 2019_TS |
| 7_68 | NAM line | Jandaroi/Fastoz10 | 2017_RW, 2017_WW, 2019_TS |
| 7_73 | NAM line | Jandaroi/Fastoz10 | 2017_RW, 2017_WW, 2019_TS |
| 7_82 | NAM line | Jandaroi/Fastoz10 | 2017_RW, 2017_WW, 2019_TS |
| 7_99 | NAM line | Jandaroi/Fastoz10 | 2017_RW, 2017_WW, 2019_TS |
| 8_101 | NAM line | Jandaroi/Fastoz6 | 2017_RW, 2017_WW, 2019_TS |
| 8_103 | NAM line | Jandaroi/Fastoz6 | 2017_RW, 2017_WW, 2019_TS |
| 8_13 | NAM line | Jandaroi/Fastoz6 | 2017_RW, 2017_WW, 2019_TS |
| 8_130 | NAM line | Jandaroi/Fastoz6 | 2017_RW, 2017_WW, 2019_TS |
| 8_143 | NAM line | Jandaroi/Fastoz6 | 2017_RW, 2017_WW |
| 8_145 | NAM line | Jandaroi/Fastoz6 | 2017_RW, 2017_WW |
| 8_163 | NAM line | Jandaroi/Fastoz6 | 2017_RW, 2017_WW |
| 8_17 | NAM line | Jandaroi/Fastoz6 | 2017_RW, 2017_WW, 2019_TS |
| 8_170 | NAM line | Jandaroi/Fastoz6 | 2017_RW, 2017_WW, 2019_TS |
| 8_176 | NAM line | Jandaroi/Fastoz6 | 2017_RW, 2017_WW |
| 8_179 | NAM line | Jandaroi/Fastoz6 | 2017_RW, 2017_WW |
| 8_43 | NAM line | Jandaroi/Fastoz6 | 2017_RW, 2017_WW, 2019_TS |
| 8_45 | NAM line | Jandaroi/Fastoz6 | 2017_RW, 2017_WW, 2019_TS |
| 8_50 | NAM line | Jandaroi/Fastoz6 | 2017_RW, 2017_WW, 2019_TS |
| 8_61 | NAM line | Jandaroi/Fastoz6 | 2017_RW, 2017_WW, 2019_TS |
| 8_76 | NAM line | Jandaroi/Fastoz6 | 2017_RW, 2017_WW, 2019_TS |
| 8_79 | NAM line | Jandaroi/Fastoz6 | 2017_RW, 2017_WW, 2019_TS |
| 8_91 | NAM line | Jandaroi/Fastoz6 | 2017_RW, 2017_WW, 2019_TS |
| 9_121 | NAM line | Jandaroi/Fastoz2 | 2017_RW, 2017_WW, 2019_TS |
| 9_13 | NAM line | Jandaroi/Fastoz2 | 2017_RW, 2017_WW, 2019_TS |
| 9_34 | NAM line | Jandaroi/Fastoz2 | 2017_RW, 2017_WW, 2019_TS |
| 9_40 | NAM line | Jandaroi/Fastoz2 | 2017_RW, 2017_WW, 2019_TS |
| 9_50 | NAM line | Jandaroi/Fastoz2 | 2017_RW, 2017_WW, 2019_TS |
| 9_55 | NAM line | Jandaroi/Fastoz2 | 2017_RW, 2017_WW, 2019_TS |

| | | | |
|---|---|---|---|
| 9_61 | NAM line | Jandaroi/Fastoz2 | 2017_RW, 2017_WW, 2019_TS |
| 9_65 | NAM line | Jandaroi/Fastoz2 | 2017_RW, 2017_WW, 2019_TS |
| 9_69 | NAM line | Jandaroi/Fastoz2 | 2017_RW, 2017_WW, 2019_TS |
| 9_7 | NAM line | Jandaroi/Fastoz2 | 2017_RW, 2017_WW, 2019_TS |
| 9_79 | NAM line | Jandaroi/Fastoz2 | 2017_RW, 2017_WW, 2019_TS |
| 9_8 | NAM line | Jandaroi/Fastoz2 | 2017_RW, 2017_WW, 2019_TS |
| 9_80 | NAM line | Jandaroi/Fastoz2 | 2017_RW, 2017_WW, 2019_TS |
| 9_83 | NAM line | Jandaroi/Fastoz2 | 2017_RW, 2017_WW, 2019_TS |
| 9_92 | NAM line | Jandaroi/Fastoz2 | 2017_RW, 2017_WW, 2019_TS |
| 9_95 | NAM line | Jandaroi/Fastoz2 | 2017_RW, 2017_WW, 2019_TS |
| 9_99 | NAM line | Jandaroi/Fastoz2 | 2017_RW, 2017_WW, 2019_TS |
| DBA Aurora | Parent | Tamaroi*2/Kalka//RH920318/Kalka///Kalka*2/Tamaroi | 2017_RW, 2017_WW, 2019_TS, 2020_WW |
| Fadda98 | Parent | Awl2/Bit = Awalbit9 | 2017_RW, 2017_WW, 2019_TS, 2020_WW |
| Fastoz10 | Parent | Younes/TdicoAlpCol//Korifla = Trouve | 2017_RW, 2017_WW, 2019_TS, 2020_WW |
| Fastoz2 | Parent | T.polonicumTurkeyIG45272/6/ICAMORTA0463/5/ Mra1/4/Aus1/3/Scar/GdoVZ579//Bit | 2017_RW, 2017_WW, 2019_TS, 2020_WW |
| Fastoz3 | Parent | Msbl1//Awl2/Bit/3/T.dicoccoidesSYRIG117887 | 2017_RW, 2017_WW, 2019_TS, 2020_WW |
| Fastoz6 | Parent | Azeghar1/6/Zna1/5/Awl1/4/Ruff//Jo/Cr/3/F9.3/7/Azeghar1//Msbl1/Quarmal | 2017_RW, 2017_WW, 2019_TS, 2020_WW |
| Fastoz7 | Parent | CandocrossH25/Ysf1//CM829/CandocrossH25 | 2017_RW, 2017_WW, 2019_TS, 2020_WW |
| Fastoz8 | Parent | MorlF38//Bcrch1/Kund1149/3/Bicrederaa1/Miki = Kunmiki | 2017_RW, 2017_WW, 2019_TS, 2020_WW |
| Jandaroi | Parent | 110780/111587 | 2017_RW, 2017_WW, 2019_TS, 2020_WW |
| Outrob4 | Parent | Ouassel-1/4/GdoVZ 512/Cit//Ruff/Fg/3/Pin/Gre//Trob = Fadda98 | 2017_RW, 2017_WW, 2019_TS, 2020_WW |

Supplemental Table S2. Summary of results from association mapping of days to flowering (DTF), plant height (PH) and spike length (SL) in the 2020 field trial.

| Trait | SNP | Chr | MAF | Pos.St (bp) | Pos.End (bp) | -log10 ($p$) | -log10($p_{\text{-FDR}}$) |
|---|---|---|---|---|---|---|---|
| DTF | 3940069 | 1B | 0.08 | 177585327 | 177585370 | 12.51 | 8.80 |
| | 1006247 | 2A | 0.35 | 36347964 | 36348032 | 4.78 | 1.37 |
| | 1022158 | 2A | 0.11 | 36347436 | 36347498 | 20.51 | 16.79 |
| | 1089102 | 2A | 0.08 | 588499216 | 588499284 | 5.27 | 2.03 |
| | 1128705 | 2A | 0.44 | 47327338 | 47327396 | 5.50 | 1.79 |
| | 2252351 | 2A | 0.40 | 35846102 | 35846170 | 5.21 | 1.62 |
| | 2256343 | 2A | 0.11 | 36364298 | 36364366 | 17.50 | 13.78 |
| | 3021936 | 2A | 0.29 | 42408208 | 42408245 | 10.69 | 6.97 |
| | 3949957 | 2A | 0.25 | 93271740 | 93271808 | 5.32 | 1.93 |
| | 4540014 | 2A | 0.10 | 58991306 | 58991340 | 4.70 | 1.50 |
| | 12779268 | 2A | 0.17 | 42489135 | 42489203 | 14.02 | 10.31 |
| | 1108975 | 2B | 0.07 | 55930502 | 55930570 | 11.87 | 8.16 |
| | 1379165 | 2B | 0.19 | 68574778 | 68574834 | 9.88 | 6.17 |
| | 3028459 | 2B | 0.06 | 54217192 | 54217234 | 15.76 | 12.05 |
| | 3064800 | 2B | 0.40 | 53972352 | 53972418 | 7.54 | 3.82 |
| | 3939176 | 2B | 0.28 | 560198210 | 560198238 | 5.13 | 1.71 |
| | 1130263 | 5A | 0.06 | 417840038 | 417840097 | 14.92 | 11.20 |
| | 1038214 | 6A | 0.49 | 595044480 | 595044548 | 5.02 | 1.60 |
| | 1765417 | 6A | 0.07 | 492965420 | 492965470 | 4.61 | 1.37 |
| | 1865002 | 6A | 0.41 | 556438754 | 556438822 | 4.72 | 1.46 |
| | 3027755 | 6A | 0.07 | 493520408 | 493520447 | 5.03 | 1.62 |
| | 1083712 | 6B | 0.11 | 513202003 | 513202071 | 4.36 | 1.30 |

| | | | | | | | |
|---|---|---|---|---|---|---|---|
| | 1106155 | 6B | 0.20 | 407590 | 407658 | 4.39 | 1.37 |
| | 1127634 | 6B | 0.50 | 677168237 | 677168304 | 5.19 | 1.77 |
| | 1137402 | 7A | 0.12 | 160972788 | 160972839 | 5.17 | 1.93 |
| | 1274875 | 7A | 0.13 | 85270676 | 85270739 | 5.35 | 1.93 |
| | 3033959 | 7A | 0.16 | 634889009 | 634888961 | 5.39 | 2.03 |
| | 3064816 | 7A | 0.16 | 634166310 | 634166378 | 4.61 | 1.50 |
| | 4440397 | 7A | 0.16 | 635288063 | 635288131 | 4.32 | 1.30 |
| | 1013473 | 7B | 0.06 | 680262333 | 680262393 | 5.00 | 1.88 |
| | 1099843 | 7B | 0.19 | 650755841 | 650755909 | 4.88 | 1.77 |
| | 1713765 | 7B | 0.07 | 621873456 | 621873524 | 4.70 | 1.46 |
| | 7354393 | 7B | 0.05 | 606979809 | 606979841 | 4.39 | 1.30 |
| PH | 4009205 | 2A | 0.27 | 982116 | 982183 | 4.73 | 1.54 |
| | 1698984 | 2A | 0.22 | 131154183 | 131154248 | 4.5 | 1.54 |
| | 1017668 | 2A | 0.17 | 695473023 | 695473091 | 4.5 | 1.54 |
| | 1088708 | 4B | 0.5 | 637884852 | 637884784 | 5.3 | 1.58 |
| | 3064427 | 5B | 0.18 | 533724323 | 533724367 | 4.8 | 1.54 |
| | 5411254 | 7A | 0.2 | 32647340 | 32647399 | 4.48 | 1.54 |
| SL | 1215020 | 1B | 0.47 | 640187467 | 640187527 | 4.65 | 1.78 |
| | 4992547 | 3A | 0.22 | 618058814 | 618058882 | 5.86 | 2.75 |
| | 1091678 | 4A | 0.46 | 636321605 | 636321665 | 8.84 | 5.13 |
| | 3954609 | 4A | 0.12 | 190565472 | 190565511 | 4.67 | 1.78 |
| | 1055097 | 5A | 0.09 | 639650884 | 639650944 | 6.51 | 3.27 |
| | 982085 | 5A | 0.46 | 43161899 | 43161953 | 4.79 | 1.78 |
| | 1092206 | 6A | 0.39 | 543489106 | 543489174 | 7.16 | 3.75 |

Chr, chromosome; MAF, minor allele frequency; Pos.St, the start of the SNP position in base pair (bp) on the 'Svevo' durum reference genome; Pos.End, the end of the SNP position on the 'Svevo' durum reference genome; -log10 (*p*), -log10 of uncorrected *p* value of marker-trait association; -log10($p_{-FDR}$), -log10 of *p* value adjusted by false discovery rate.
